# Expression study of Krabbe Disease *GALC* missense variants – Insights from quantification profiles of residual enzyme activity, secretion and psychosine levels

**DOI:** 10.1101/2024.10.17.618938

**Authors:** Hui Peng, Ying-Wai Lam, Zitao Zhou, Aimee R. Herdt, Michael H. Gelb, Chris W. Lee

**Affiliations:** Biomedical Research Institute of New Jersey (BRInj), Cedar Knolls, NJ, USA; Atlantic Health System, Morristown, NJ, USA; MidAtlantic Neonatology Associates (MANA), Morristown, NJ, USA; Department of Biology & Vermont Biomedical Research Network Proteomics Facility, University of Vermont, Burlington, Vermont, USA; Departments of Chemistry and Biochemistry, University of Washington, Seattle, WA, USA

**Keywords:** Globoid cell leukodystrophy, Krabbe disease, GALC, galactosylceramidase, galactosylsphingosine, psychosine, lysosomal storage disorder, genotype-phenotype correlation, missense mutation variant, expression study, protein trafficking, secretion, lysosome, crispr-cas9 knockout cell

## Abstract

Krabbe disease (KD) is an autosomal recessive lysosomal storage disorder caused by loss-of-function mutations in the *GALC* gene, which encodes for the enzyme galactosylceramidase (GALC). GALC is crucial for myelin metabolism. Functional deficiency of GALC leads to toxic accumulation of psychosine, dysfunction and death of oligodendrocytes, and eventual brain demyelination. To date, 46 clinically-relevant, pathogenic GALC missense mutations (MMs) have been identified in KD patients. These MMs are present in ∼70% of KD cases reported over 8 published studies between 1996 – 2019. However, the mechanisms by which these MMs lead to GALC functional deficiency and their correlations with clinical phenotype remain poorly understood.

To address this, we generated a *GALC*-knockout human oligodendrocytic cell line (MO3.13/ *GALC*-KO) using CRISPR-Cas9 method to assess GALC function and GALC secretion. We evaluated 5 polymorphic and 31 clinically-relevant MM variants (MMVs) using transient expression assays. Our results showed that 26 MMVs, including 10 co-variants with p.I562T, reduced GALC activity by 92% - 100% compared to wild-type GALC (WT-GALC). MMVs from infantile-onset KD patients produced < 2% of WT activity, whereas those associated with juvenile- and adult-onset cases retained up to 7% of WT activity. Residual GALC activity was correlated with mature, lysosomal GALC protein levels (Pearson r = 0.93, P<0.0001). Many low-activity MMVs did not correspondingly impair GALC secretion. Twenty-one of the 26 low-activity MMVs showed a 21% - 100% reduction in sec-GALC levels, indicating varying degrees of GALC mis-trafficking among these variants.

Importantly, GALC activity among MMVs strongly correlates with clinical disease severity, based on the age of symptom onset in patients with either homozygous MM (Pearson r = 0.98, P<0.0001, n = 7) or compound heterozygous (Pearson r = 0.94, P<0.0001, n = 12) MM-null mutation genotypes. Thus, our data suggests that GALC activity could serve as a prognostic disease indicator under specific experimental conditions. We further investigated the impact of pathogenic MMVs on psychosine accumulation, a key biomarker for KD. Psychosine levels were 21-fold higher in mock control cells compared to WT-GALC transfected cells (mock = 0.349 pmol/mg, WT-GALC = 0.016 pmol/mg), but negatively correlated with GALC activity among pathogenic MMVs (Pearson r = −0.63, P < 0.01, n = 15). Although psychosine levels were higher in most MMVs associated with infantile-onset KD, no significant correlations with clinical onset were detected.

Overall, our study provides a comprehensive quantitative analysis of the functional deficits and mis-trafficking associated with clinically-relevant GALC MMVs, enhancing our understanding of the molecular genetics and genotype-phenotype correlations of the *GALC* gene in Krabbe disease.

## Introduction

Globoid cell leukodystrophy or Krabbe disease (KD) (OMIM 245200) is an inherited demyelinating disorder caused by a deficiency of galactosylceramidase (GALC; EC 3.2.1.46). GALC function is essential for myelin metabolism^1–5^. The loss of GALC activity leads to the accumulation of psychosine, as well as the dysfunction and death of oligodendrocytes and Schwann cells; thus leading to demyelination in both the central and peripheral nervous systems (CNS and PNS) ^6–9^.

To date, 243 KD-related mutations in the *GALC* gene have been classified as pathogenic by the American College of Medical Genetics and Genomics (ACMG) guidelines^10^ in the NCBI ClinVar archive^11^. Among those, 46 are missense mutation variants (MMVs). Statistics from published studies between 1996 and 2019 document KD patients from different geographical regions with varying disease severity and show that 38% to 95% carried at least one *GALC* MMV allele (Table 1)^12–19^. In the Bascou et al. study, 19 of 20 patients had at least one *GALC* MMV allele and 12 had infantile-onset KD^13^. These observations highlight that *GALC* MMVs are common in the KD population.

**Table 1.** Frequency of *GALC* missense mutation alleles in KD population.

| Study | KD Patients |  | MM Allele | Age-of-onset |  | Patient's Geographic |
| --- | --- | --- | --- | --- | --- | --- |
|  | Case | MM Allele | Frequency | Range (month) | Median (month) |  |
| Madsen et al. 2019 <sup>12</sup> | 29 | 11 | 38% | 1 - 36 | 3.3 | Denmark |
| Bascou et al. 2018 <sup>13</sup> | 20 | 19 | 95% | 6 - 36 | 9 | United States |
| Duffner et al. 2012 <sup>14</sup> | 14 | 13 | 93% | 8 - 384 | 15 | European |
| Duffner et al. 2011 <sup>15</sup> | 16 | 7 | 44% | 2 - 5 | 3 | European |
| Tappino et al. 2010 <sup>16</sup> | 30 | 21 | 70% | 1 - 312 | 5 | Italian |
| Xu et al. 2006 <sup>17</sup> | 14 | 12 | 86% | 4 - 828 | 16 | Japan |
| Fu et al. 1999 <sup>18</sup> | 8 | 6 | 75% | Infantile | N.A. | Japan |
| De Gasperi et al. 1996 <sup>19</sup> | 10 | 8 | 80% | 5 - 216 | 54 | United States |

Case: Total number of KD patients reported
MM Allele: Number of patients with at least one *GALC* missense mutation allele
Frequency: Percentage of patients with at least one MM allele

KD is an autosomal recessive disorder, such that most KD patients are compound heterozygous for *GALC* mutations. Patients of northern European descent most frequently carry a 30 kb deletion on one allele^20, 21^ and a MMV on the other allele^14, 15^. Null mutations, such as the 30 kb deletion, eliminate normal *GALC* transcripts and GALC protein, leading to complete functional deficiency. By contrast, MMVs impair GALC function via alternative mechanisms, such as protein misfolding, instability, mis-trafficking and catalytic inactivation.

GALC is a glycoprotein that traffics to the lysosome through the mannose 6-phosphate receptor-mediated pathway^22^. Briefly, GALC is modified post-translationally at Golgi. Precursor GALC (pre-GALC) is then secreted out of the cell by secretory vesicles to become secreted GALC (sec-GALC) or transported to lysosomes via the endosomal pathway (Figure 1A). Of note, upon delivery to the lysosome, GALC undergoes a unique proteolytic cleavage event that generates amino-terminal (∼50 kDa) and carboxyl-terminal (C-terminal, ∼30 KDa) fragments^23^, held together by disulfide bridges^24^. Concomitantly, the extreme C-terminal end of the 30 kDa fragment is proteolytically trimmed further, rendering the fragment undetectable using antibodies against protein tags fused to the C-terminal end^25, 26^ (Figure 1B). Although the functional effects of the proteolytic cleavages have not been determined, we expect levels of cleaved GALC fragment (lys-GALC) to be proportional to the mature, active form of GALC in lysosomes^25–27^.

**Figure 1.**
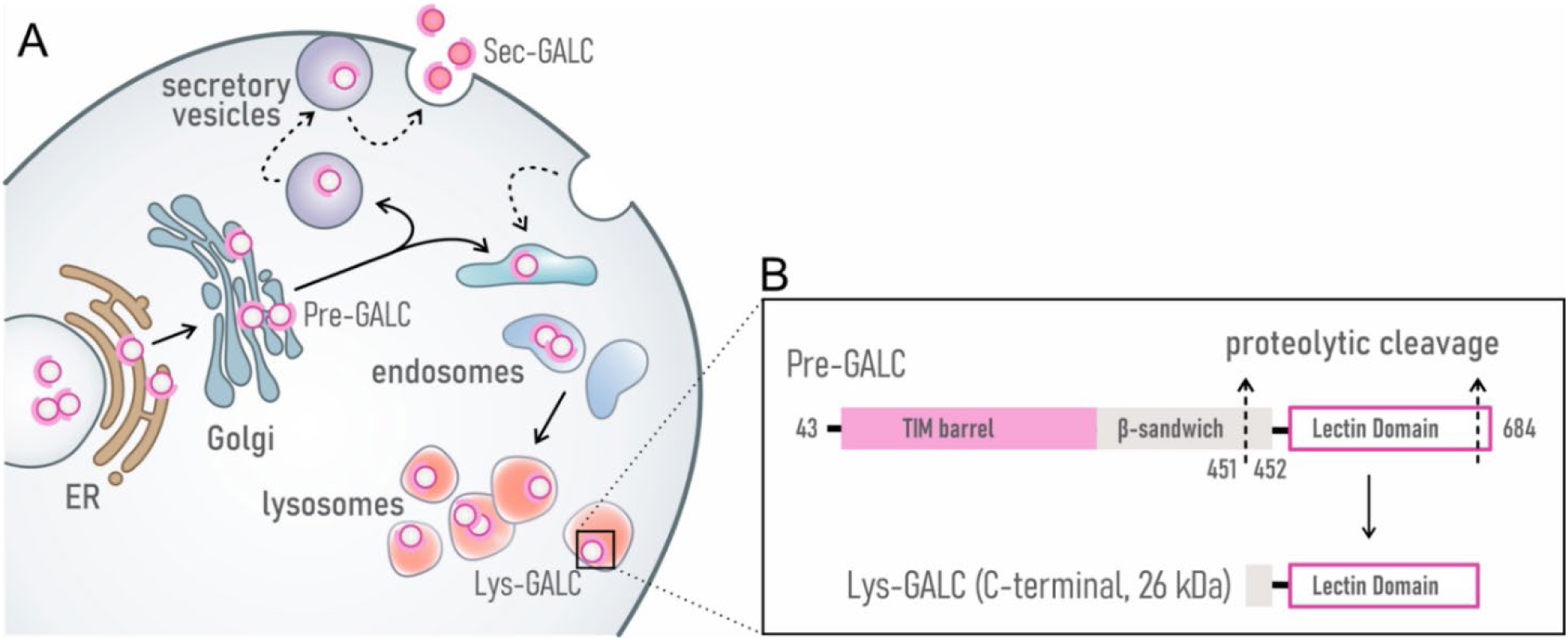
Subcellular trafficking and lysosomal processing of GALC protein. (A) Schematic illustration of GALC trafficking through subcellular compartments. After protein synthesis in the ER, precursor GALC (Pre-GALC) is transported through the Golgi to 1) the extracellular space via secretory vesicles, where it forms secreted GALC (Sec-GALC), or 2) lysosomes via the endosomal pathway, where it becomes lysosomal GALC (Lys-GALC). (B) Lysosomal processing of GALC. Pre-GALC undergoes proteolytic cleavage at sites located in the β-sandwich and lectin domains to form mature GALC within lysosomes. A carboxyl-terminal lys-GALC fragment can be detected by Western blot using the CL13.1 anti-GALC antibody.

GALC MMVs are evenly distributed among the three protein domains, the triosephosphate isomerase (TIM) barrel domain (57 – 353 aa), β-sandwich domain (354 – 468 aa) and lectin binding domain (488 – 684 aa) (Figure 2)^24^. It has been shown that p.T529M, p.L634S and p.L645R induce GALC mis-trafficking and reduce lysosomal localization and secretion^25–27^. Missense mutation at the p.R396 residue, which is essential for substrate binding^28^, impairs GALC activity^29, 30^, but not trafficking of the GALC mutant protein^27^. These studies provide mechanistic insight into a few pathogenic MMVs; however, the question of how most other variants contribute to pathogenicity remains.

**Figure 2.**
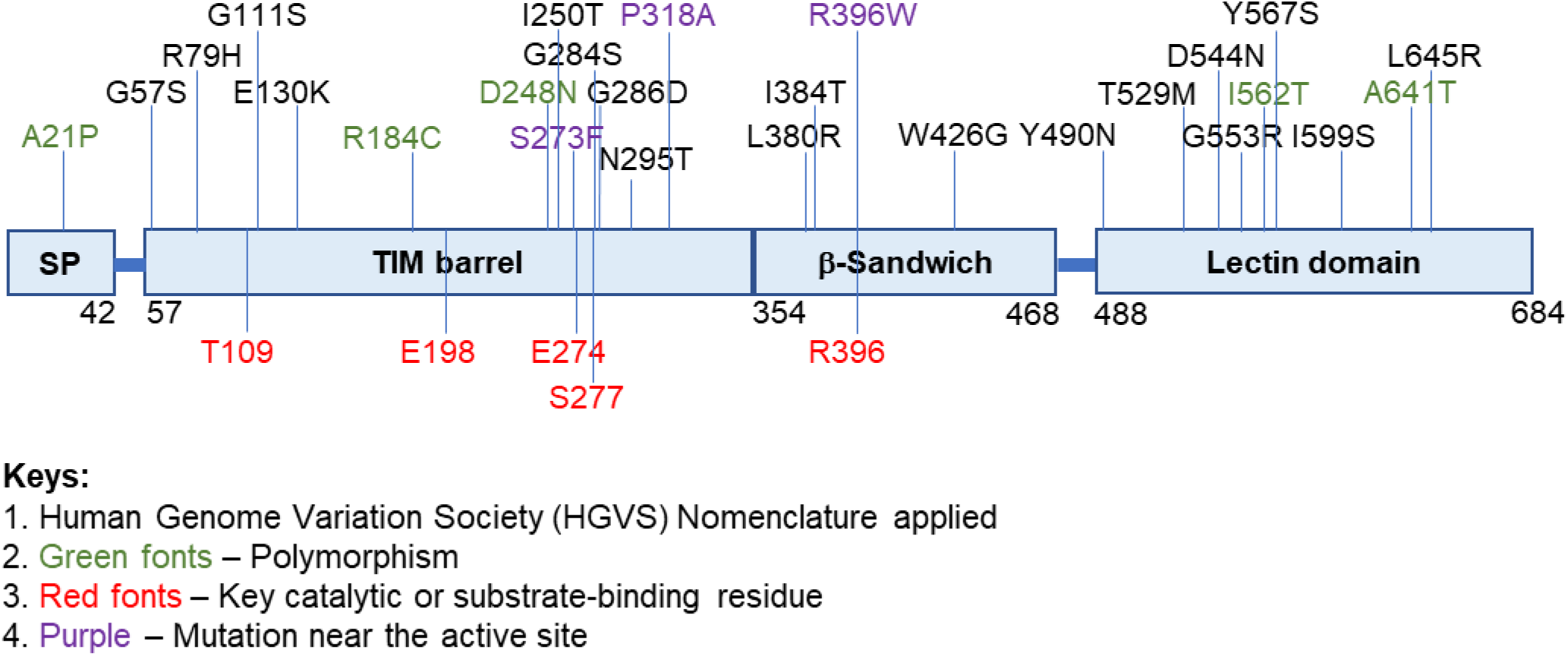
Locational distribution of missense mutation variants on the human GALC protein. The schematic diagram shows that distribution of clinically-relevant missense mutation variants (MMVs), common polymorphic variants (green) and substrate binding site residues (red) on the GALC protein. The main structural domains of GALC are indicated: signal peptide (SP), TIM barrel domain, β-sandwich domain and lectin-binding domain. MMVs predicted to affect substrate binding are labeled purple color. Potential N-glycosylation sites are marked with asterisks.

To understand how individual *GALC* MMVs contribute to pathogenicity in KD, we selected 21 pathogenic MMVs and characterized their effect on GALC activity and GALC trafficking. We also examined ten *GALC* MMVs previously found to occur *in cis* with p.I562T, a known pseudo-deficiency variant frequently identified as an allelic background with other variants^16, 31^. Some MMVs, including p.S273F, p.P318A and p.R396W, are likely to alter GALC’s active site, and therefore, disrupt catalytic function^24^. Two MMVs, p.N295T and p.D544N, previously shown to alter N-glycosylation status of GALC proteins^25, 27^, were also included in the MMV panel. Altogether, we designed a systematic expression study of 5 polymorphic variants and 31 clinically-relevant MMVs, including 21 single- and 10 p.I562T co-variants, in human oligodendrocytic MO3.13 cells. The list of *GALC* MMVs examined in this study and related information are shown in Table 2. To eliminate the background signal from endogenous GALC protein and maximize sensitivity, a Crispr-Cas9 method was used to generate *GALC* knockout cells. This strategy is crucial for MMVs with low GALC activity overlapping with endogenous levels. Residual GALC activity, lys-GALC protein levels, pre-GALC protein levels, sec-GALC protein levels and psychosine levels were examined in a quantitative manner, in order to understand the molecular effect elicited by these MMVs on GALC function and trafficking, as well as to establish correlations to KD clinical severity.

**Table 2.** Clinically-relevant GALC MMVs examined in this study.

| **GALC Variant** | **Polymorphic MM** | ***In cis* w/ p.I562T** | **Patient Geographic** | **Note** | **Reference** |
| --- | --- | --- | --- | --- | --- |
|  | p.A641T |  |  |  |  |
|  | p.A21P |  |  |  |  |
|  | p.R184C |  |  | *In cis* with 30 kb deletion | 21 |
|  | p.D248N |  |  |  |  |
|  | p.I562T |  |  | Common pseudo-deficiency variant | 31 |
| **WT** |  |  |  |  |  |
| **p.G57S** |  | Yes | Italian |  | 13, 14, 47 |
| **p.R79H** |  | Yes | Italian |  | 16, 19 |
| **p.G111S** |  |  |  |  | 19 |
| **p.E130K** |  |  | Italian | Present in the Twi-5J mouse model | 16, 47, 62 |
| **p.I250T** |  |  | Greek |  | 17, 19, 29 |
| **p.S273F** |  |  | Japanese | Near SBS | 17 |
| **p.G284S** |  | Yes |  |  | 13, 14, 17, 19 |
| **p.G286D** |  | Yes | Japanese; European | Late-onset KD mutation | 14, 16, 17, 19, 29 |
| **p.N295T** |  |  | Italian |  | 16 |
| **p.P318A** |  |  | Japanese | Near SBS | 17 |
| **p.L380R** |  |  | Japanese |  | 17 |
| **p.I384T** |  | Yes | Italian |  | 16 |
| **p.R396W** |  |  |  | Severely affect substrate binding | 12, 15, 16 |
| **p.W426G** |  | Yes |  |  | 18 |
| **p.Y490N** |  | Yes | Italian |  | 15, 16 |
| **p.T529M** |  | Yes | Northern European | High frequency | 12-15 |
| **p.D544N** |  | Yes | Israeli Arab | Hyper-glycosylated | 25, 63, 64 |
| **p.G553R** |  |  | Italian |  | 16 |
| **p.Y567S** |  | Yes | Northern European | High frequency | 12-16 |
| **p.I599S** |  |  | Israeli Druze |  | 63 |
| **p.L645R** |  |  |  |  | 13, 65 |

*In cis* w/ p.I562T: MMV is frequently inherited in-cis with the p.I562T background
SBS: Substrate-binding site

## Materials and Methods

### MO3.13/ *GALC*-KO cell line

Guided RNA targeting sequence 1 (sgRNA-GALC1: 5’ GAAAGCTATGACTGCGGCCG 3’) and 2 (sgRNA-GALC2: 5’ TACGTGCTCGACGACTCCGA 3’) were subcloned individually into pX459 v2.0 (gift from Dr. Feng Zhang, Addgene plasmid #62988^32^). GALC targeting sequences were further tested by the Off-Spotter software to minimize potential off target effect. Vectors were co-transfected into MO3.13 cells (Cedarlanes, Burlington, NC) with puromycin selection for 3 days. After 4 weeks of colony formation, multiple colonies were isolated and expanded for sequence validation. The sgRNAs were designed to specifically target 2 protospacer adjacent motif (PAM) sites on exon 1 of the *GALC* gene. Positive clones with cleavage on the target sites created an 85-bp deletion and a downstream nonsense mutation at 319 nt of exon 2 (Figure 2A and 2B). We confirmed the absence of GALC protein (Figure 2C) and GALC activity (Figure 2D) in the *GALO*-KO cell line by western blot and GALC activity assay, respectively. Substrate accumulation in the *GALC*-KO cell line was confirmed by psychosine analysis (Figure 2E).

### Generation of *GALC* wildtype and mutant expression constructs

Full-length, human, wild-type *GALC* cDNA (SC120078, NM_000153) was originally purchased from Origene (Rockville, MD). The cDNA was subcloned into the pBApo-CMV pur expression vector at the XbaI and PstI sites (Takara Bio, San Jose, CA). V5 and 6-histidine tags were included on the carboxyl-terminal for protein detection and purification purposes^25^. Point mutations were introduced to the pBApo-GALC-VH construct with the Q5 site-directed mutagenesis kit (New England Biolabs, Ipswich, MA). Non-overlapping primer pairs for each MM are listed in Supplementary Table 1. DNA sequencing was performed on all clones to ensure no random mutations were introduced and that the desired mutation was present.

### Transient GALC variants expression study

GALC expression constructs were transfected to MO3.13/ *GALC*-KO cells using the Lipofectamine 3000 reagent according to the manufacturer’s instructions (Thermofisher Scientific, Waltham, MA). Briefly, 500 ng of pBApo-GALC-VH plasmid was transfected to pre-plated cells on 12-well culture plates at 70%-80% density. The cells were cultured in completed medium (Dulbecco’s modified eagle medium with 10% (v/v) fetal bovine serum, and 1% (v/v) penicillin-streptomycin) for 72 hours. To account for any variability in transfection efficiency, GALC activity and protein levels were normalized to puromycine N-acetyltransferase, which was expressed by the same DNA construct. Results presented in Table 3 are mean values ± standard error mean (SEM) obtained from four independent experiments.

**Table 3.**
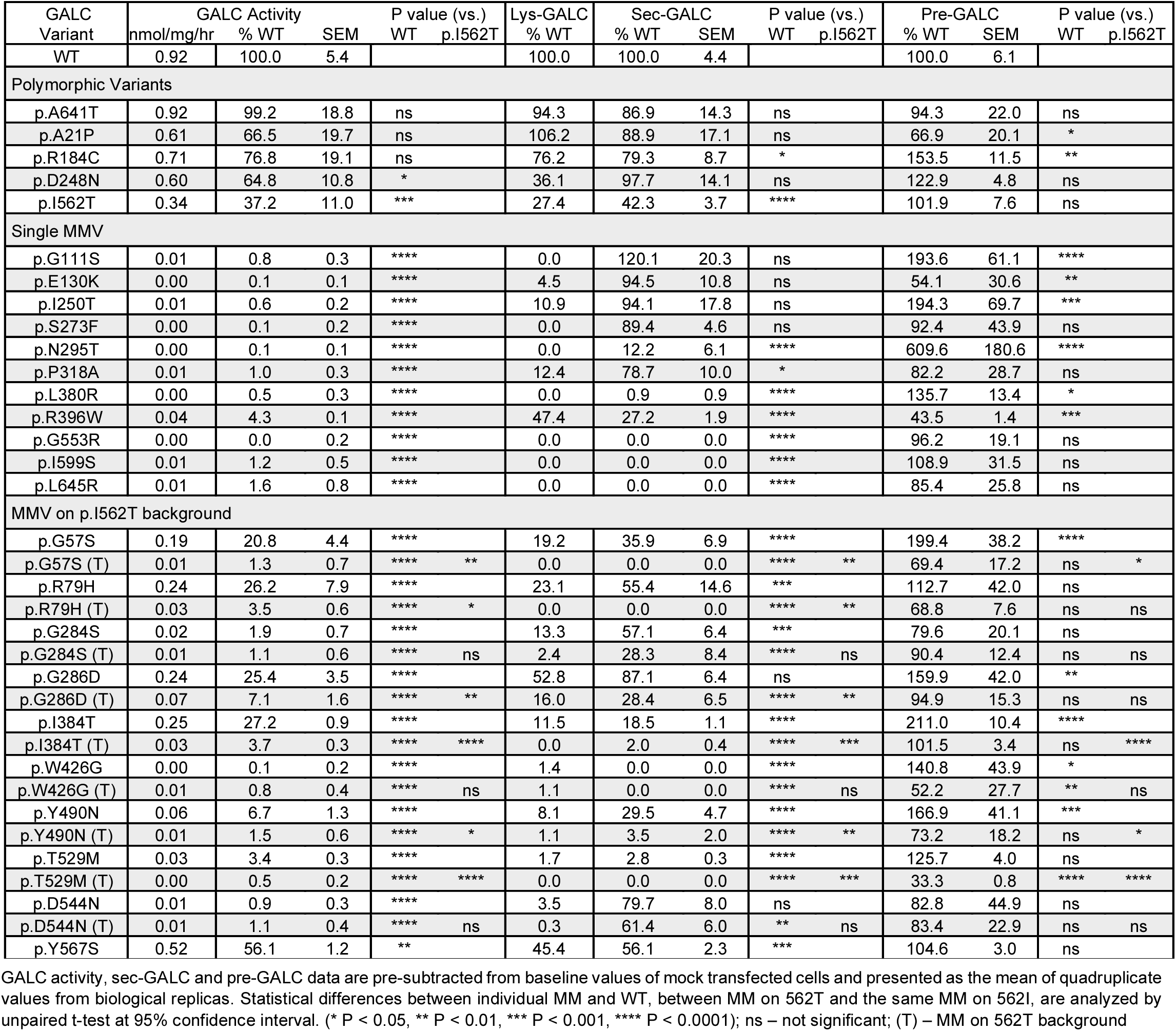
Molecular characterization of GALC MMVs in MO3.13/ *GALC*-KO transiently-expressed cell models.

### Cell harvesting and sample preparation for analysis

At harvesting, culture medium was collected and centrifuged to remove floating cells and debris. The medium supernatant was saved for analysis of sec-GALC by ELISA. Cells were washed with cold PBS after removal of culture medium. Residual PBS was removed by vacuum suction. Cells were lysed using the M-Per mammalian extraction reagent supplemented with the 1x Halt protease inhibitor cocktail (Thermo Fisher Scientific, Waltham, MA) on ice for 20 minutes. The insoluble fraction was removed by centrifugation at 13,000 rcf for 30 minutes. Total soluble lysate was transferred to a clean tube and stored on ice. Protein concentration of lysate was measured by Pierce BCA protein assay (Thermo Fisher Scientific, Waltham, MA). Cell lysate and medium supernatant were kept cold, but never frozen, prior to GALC activity, lys-GALC, pre-GALC and sec-GALC analysis.

### In vitro GALC activity assay

Residual GALC enzymatic activity in cell lysate was measured by a fluorescence substrate turnover assay ^33^. Briefly, about 20 μg total protein was added to a reaction cocktail containing 0.1M citric acid, 0.2M sodium phosphate, 7 mg/ml sodium taurocholate, 2 mg/ml oleic acid and 0.5 mg/ml 6-hexadecanoylamino-4-methylumbelliferyl-β-D-galactopyranoside (HMGal; Biosynth Inc., Newbury, U.K.) at pH 4.5. The reaction was incubated for 4 hours at 37°C and stopped by adding 2 volumes of 0.1M glycine/ 0.1M sodium hydroxide solution and 4 volumes of absolute ethanol. The concentration of reaction product was determined by fluorescence plate reader (ex 385nm/ em 450nm). A 4-methylumbelliferone standard curve was ran in parallel for calculation of absolute GALC activity unit expressed in nmol per hour per milligram protein.

### Western blot

Cultured cells were lysed in M-Per lysis buffer (ThermoFisher Scientific) supplemented with protease inhibitors. After centrifugation, the protein concentration of the supernatant was determined by BCA assay for normalization, and 30 μg of total protein was resolved by SDS-PAGE with 10-20% Tris-glycine gel. Gel proteins were transferred to PVDF membranes by Criterion blotter (Bio-Rad, Hercules, CA) according to manufacturer’s instructions. The protein blot was blocked in 5% non-fat milk for 1 hour at room temperature, followed by incubation with primary antibody for 16-18 hours at 4°C. The blot was probed with HRP-conjugated secondary antibody for 1 hour at room temperature, followed by signal development using the Immobilon Western Chemiluminescent HRP substrate (MilliporeSigma). WB images were obtained using Image-quant LAS 4000. Primary antibodies for WB analysis include anti-GALC (CL13.1, 1:4000) ^34^ and anti-GAPDH (1:100,000, Proteintech, Rosemount, IL).

### GALC enzyme-linked immunosorbent assay (ELISA)

Secretory GALC levels were measured using 50 μL of culture supernatant added to a Nunc Maxisorp^TM^ 96-well plate pre-coated with 2 μg/ml anti-V5 tag antibody and pre-blocked in complete growth medium (DMEM with 10% FBS). Standard curve samples were prepared by diluting recombinant human GALC protein (R&D Systems) in culture supernatant from KO cells to 0.8, 1.6, 3.2, 6.3, 12.5, 25 and 50 ng/ml. Samples and standards were incubated for 2 hours. Anti-GALC antibody, CL13.1, followed by anti-mouse IgG1-HRP secondary antibody, both diluted in sample buffer (PBS with 1% BSA) to 0.5 μg/ml, were added to the plate and incubated for an hour. TMB substrate was added to the plate to develop the signal and incubated for 10 - 30 min. Diluted sulfuric acid (0.16 M) was added to stop the reaction. The plate was read on the SpectraMax Plus 384 microplate reader (450 nm). All procedures except antibody coating were performed at room temperature. Plates were washed between blocking, sample addition and antibody addition using PBS with 0.05% Tween-20. GALC protein concentration was calculated by interpolation of absorbance values to a normalized standard curve.

Intracellular pre-GALC levels were measured as described above, except 30 μg of protein from total lysate was added to pre-coated ELISA plates and pre-blocked in sample buffer. Standard curve samples were prepared by diluting recombinant GALC protein in sample buffer to the same concentration range described above.

### Psychosine measurements

Cultured cells were washed, then scrapped off in cold PBS. Cell pellets were collected by centrifugation at 13,000 rcf for 10 minutes and stored at −80 °C. Replicated set of cell pellets was lysed in M-Per lysis buffer to determine the total protein concentration for calculation of the psychosine level in pico-mole per milligrams protein. For psychosine analysis in the correlation study (Figure 12), cultured cells were washed in cold PBS then lysed by digitonin (0.1%) containing Tris-buffered saline supplemented with 1x Halt protease inhibitor cocktail. After centrifugation, the insoluble lipid pellet was saved and stored at −80 °C for analysis. The soluble lysate was analyzed for total protein concentation and residual GALC activity.

Frozen samples were sent to Gelb’s lab for psychosine analysis. Briefly, 50 μl of 0.9% sodium chloride solution was added to each pellet to homogenize. Internal standard solution (250 μl, 1 nM d5-psychosine) was then mixed and incubated with the sample for 2 hours. The mixture was centrifuged at 13000 rcf for 5 minutes. Supernatant went through SPE cleanup and was dried by speedvac. The sample was reconstituted with 100 ml of mobile phase B solution, followed by injection to the MedChem Xevo TQS system. Detailed procedures for UPLC-MS/MS analysis were recently described ^35^.

### Statistical analysis

Figure 3: Statistical differences in GALC activity (n=4) or psychosine level (n=3) between WT and GALC-KO cells were analyzed by unpaired t-test at 95% confidence interval. (** P < 0.01, *** P < 0.001).

**Figure 3.**
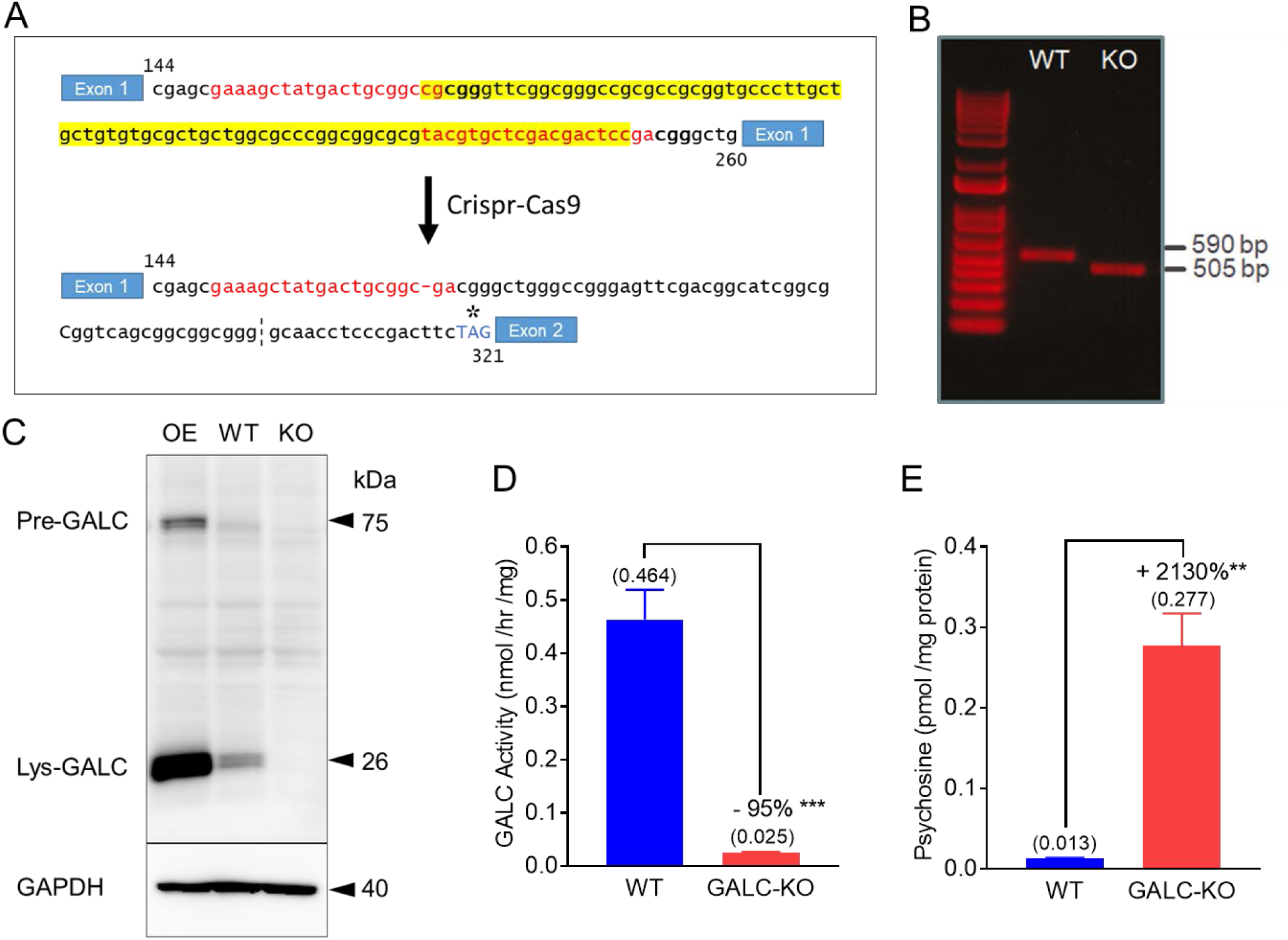
Generation and validation of MO3.13/ *GALC*-KO cells. (A) Human *GALC* gene sequences (red) adjacent to PAM sites (bolded) were targeted for Cas9-mediated cleavage by the sgRNA vectors. A downstream nonsense mutation (asterisk) is introduced upon successful targeted deletion (highlighted yellow). (B) Confirmation of targeted deletion of the 85 bp region in the KO cell line by PCR amplification, compared to control WT cells. (C) Western blot analysis of GALC and GAPDH proteins in native MO3.13 cells (WT), GALC-KO cells (KO) and GALC-overexpressing cells (OE). (D) GALC activity and (E) psychosine levels in WT and KO cells. Statistical significance is determined using an unpaired t-test (n=4, 95% confidence interval, ** P < 0.01, *** P < 0.001).

Figure 8A and 9C: Statistical differences in sec-GALC levels (n = 4) or pre-GALC levels (n = 3) among cells expressing individual MMVs vs. WT were analyzed by unpaired t-test at 95% confidence interval. (n=4, *P < 0.05, **P<0.01, ***P<0.001, ****P < 0.0001).

Figure 11: Statistical differences in (A) GALC activity, (B) sec-GALC level and (C) pre-GALC levels between cells expressing the p.562I MMV and the same MMV on the p.562T background were analyzed by unpaired t-test at 95% confidence interval. (* P < 0.05, ** P < 0.01, *** P < 0.001, **** P < 0.0001).

Table 3 and 4: Statistical differences between cells expressing individual MMVs and WT, between cells expressing MMVs on p.562T and the same MMV on the p.562I background, were analyzed by unpaired t-test at 95% confidence interval. (* P < 0.05, ** P < 0.01, *** P < 0.001, **** P < 0.0001).

Figure 6B, 6C, 7E, 8B, 8C, 9B, 10D, 10E, 11A, 11B, 11C, 12A, 12B, 12C and 12D: Correlation between groups was analyzed by Pearson correlation. Significance of the Pearson correlation coefficient (i.e., Pearson r) was determined by a P-value at 95% confidence interval. (* P < 0.05, ** P < 0.01, *** P < 0.001, **** P < 0.0001).

## Results

### *GALC* knockout background, human oligodendrocytic cell line for GALC MMVs expression study

The demyelination observed in the central nervous system of KD is caused by functional deficiency of GALC in oligodendrocytes. MO3.13 cells express various oligodendrocyte markers, including 2’,3’-cyclic-nucleotide 3’-phosphodiesterase (CNPase), galactosylceramide (GalCer) and myelin basic protein (MBP)^36, 37^, and thus, are frequently used to study molecular mechanisms related to demyelination disorders, including multiple sclerosis^38, 39^ and KD^40, 41^. Likewise, for this study, we chose to model the molecular effect of GALC MMVs in this human oligodendrocytic cell line. To prevent endogenous wildtype GALC from interfering with the measurement of exogenous GALC variants, we knocked out *GALC* gene expression using the Crispr-Cas9 genome editing method. Targeted deletion of an 85 bp nucleotide in exon 1 was confirmed by PCR amplification of genomic DNA from the *GALC*-KO cell line (Figure. 3A and 3B). The nonsense mutation likely eliminates *GALC* mRNA via the nonsense-mediated mRNA decay pathway. Absence of mature, lysosomal GALC protein (lys-GALC; 26 kDa) was confirmed by WB, compared to native MO3.13 cells with and without the over-expression of GALC (Figure 3C). Some background GALC activity was detected in the KO cells (0.025 nmol/mg/hr); however, native MO3.13 cells have 19-times higher activity (0.464 nmol/mg/hr) (Figure 3D). Psychosine levels are 21-times higher in KO cells compared to native cells (0.277 pmol/mg vs 0.013 pmol/mg, respectively); which highlights an anticipated deficiency of substrate catabolic activity (Figure 3E). Our results confirm the *GALC* knockout status of the MO3.13/ *GALC*-KO cell line.

### Differential expression of lysosomal and ceramide pathway-related proteins in the MO3.13/ GALC-KO cells

We compared the proteomic profile between the *GALC*-KO cells and native cells (Fig. 3) using isobaric tandem mass tags, a stable, isotope-based quantitative proteomic approach. A total of 4,863 proteins were identified and 2,417 quantified (Figure 4A). Three hundred and sixty-six proteins were differently expressed, (Figure 4B, p < 0.05), both upregulated (104 with KO/WT > 1.25; log2 > 0.33) and downregulated (161, KO/WT < 0.8; log2 < −0.33). Interestingly, expression of several lysosomal proteins, NPC intracellular cholesterol transporter 2 (NPC2), ganglioside GM2 activator (GM2A), N-acetylgalactosamine-6-sulfatase (GALNS) and tripeptidyl-peptidase 1 (TPP1), were reduced in the GALC-KO cells (49%, 40%, 32% and 31%, respectively). In contrast, levels of ceramide synthase 6 (CERS6) increased by 28% in GALC-KO cells. Other proteins in the *de novo* ceramide synthesis pathway, including dihydrolipoamide dehydrogenase (DLD, +49%), malate dehydrogenase (MDH2, +30%) and aldehyde dehydrogenase 2 (ALDH2, +29%), were also elevated in the *GALC*-KO cells, which suggests there is a compensatory effect of ceramide production (Figure 4C).

**Figure 4.**
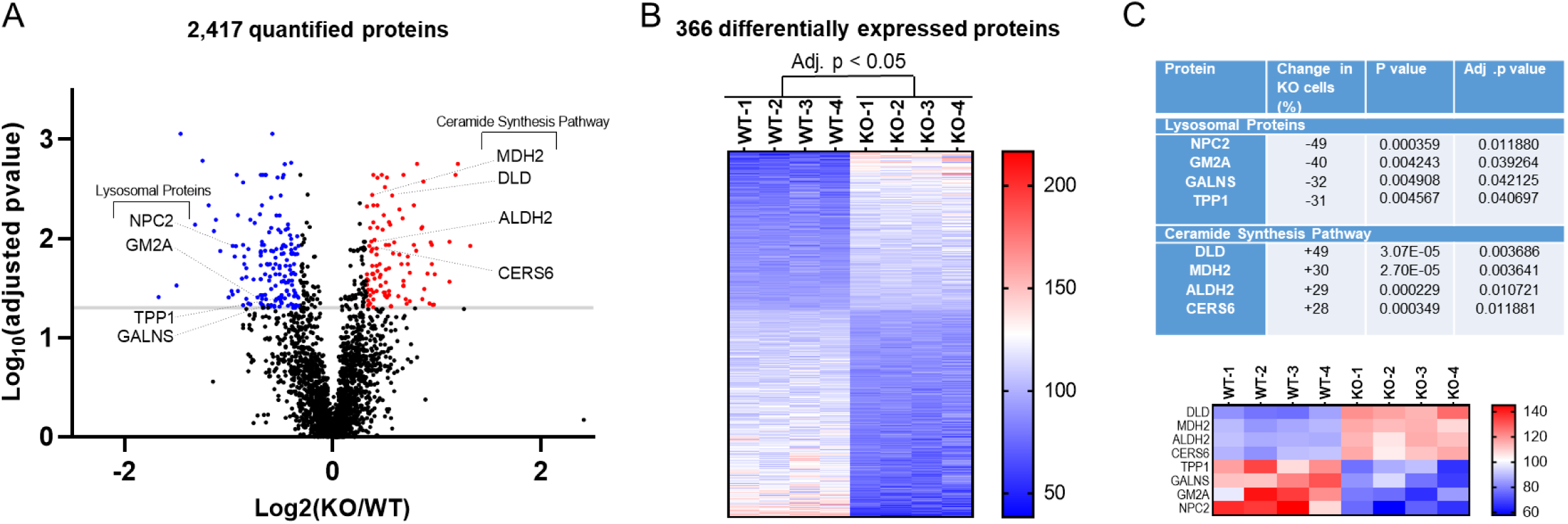
Differential proteome of MO3.13 native cells and MO3.13/ *GALC*-KO cells. (A) Proteomic profiles of MO3.13 native cells (WT) and MO3.13/ *GALC*-KO cells (KO) cells are compared. A total of 2,417 proteins were identified and quantified, and the results are displayed in a volcano plot. Three hundred and sixty-six proteins were identified as differentially expressed in KO versus WT cells (two-tailed t-test; adjusted p value (Benjamini-Hochberg) < 0.05). A P value of 0.05 (-log_10_ 0.05 = 1.301) is indicated by a grey line. Differentially expressed proteins with a 1.2-fold change (KO/WT: log_2_ < −0.33 or > 0.33) are highlighted (up regulated in red; downregulated in blue). (B) Heat map of the scaled abundances of the 366 differentially expressed proteins in descending order based on average log_2-_fold change. The color intensity scale is arbitrary (double gradient). (C) Differently expressed lysosomal proteins and proteins in the *de novo* ceramide synthesis pathway are listed. The percent change (KO vs. WT) and p values were calculated. Scaled abundances of the target proteins are shown in a heat map in descending order based on average log_2-_fold change.

### Dramatic reduction of functional activity in 20 clinically-relevant GALC MMVs

Low in vitro GALC activity in leukocytes (≤ 0.15 nmol/mg/hr) is commonly used as the first confirmatory test for KD in clinical settings ^42, 43^. Therefore, we also measured residual GALC activity in transfected cell lysate as a readout to determine the molecular effect of MMVs on GALC in MO3.13/ *GALC*-KO cells. GALC activity of the WT transfectant was two-fold greater than native MO3.13 cells (0.92 nmol/mg/hr vs. 0.46 nmol/mg/hr, respectively) (Table 3, GALC Activity). Three out of 5 polymorphic variants, p.A641T, p.A21P and p.R184C, had similar activity to WT-GALC transfected cells. The remaining 2 variants, p.D248N and p.I562T, had GALC activity reduced by 32% and 61%, respectively (Table 3; Figure 5, green bars). By contrast, 20 of the 31 KD MMVs reduced GALC activity by more than 98%, including p.G284S, p.L645R, p.Y490N(T) (p.I562T cis-background abbreviated as “(T)”), p.G57S(T), p.I599S, p.D544N(T), p.G284S(T), p.P318A, p.D544N, p.Y567S(T), p.G111S, p.W426G(T), p.I250T, p.L380R, p.T529M(T), p.S273F, p.N295T, p.E130K, p.W426G and p.G553R. Most of the MMVs known to be associated with the infantile-onset form of KD, such as p.G553R, p.T528M(T) and p.G567S(T), were found in this group (Figure 5, red bars). MMVs with activity between 2% - 7% of WT activity included variants associated with later-onset form of KD, such as p.G286D(T) and p.I384T(T) (Figure 5, orange bar). MMVs with activity between 20% and 60% are the pseudo-deficiency variant p.I562T and other single variants, including p.Y567S, p.G286D, p.I384T, p.R79H and p.G57S, likely to be non-pathogenic in the absence of the p.I562T background (Figure 5, blue bar). Our data suggests that residual GALC activity from MMVs expressed in MO3.13/ *GALC*-KO cells has the potential to be used as a readout to predict, based on percent activity, a clinical KD form.

**Figure 5.**
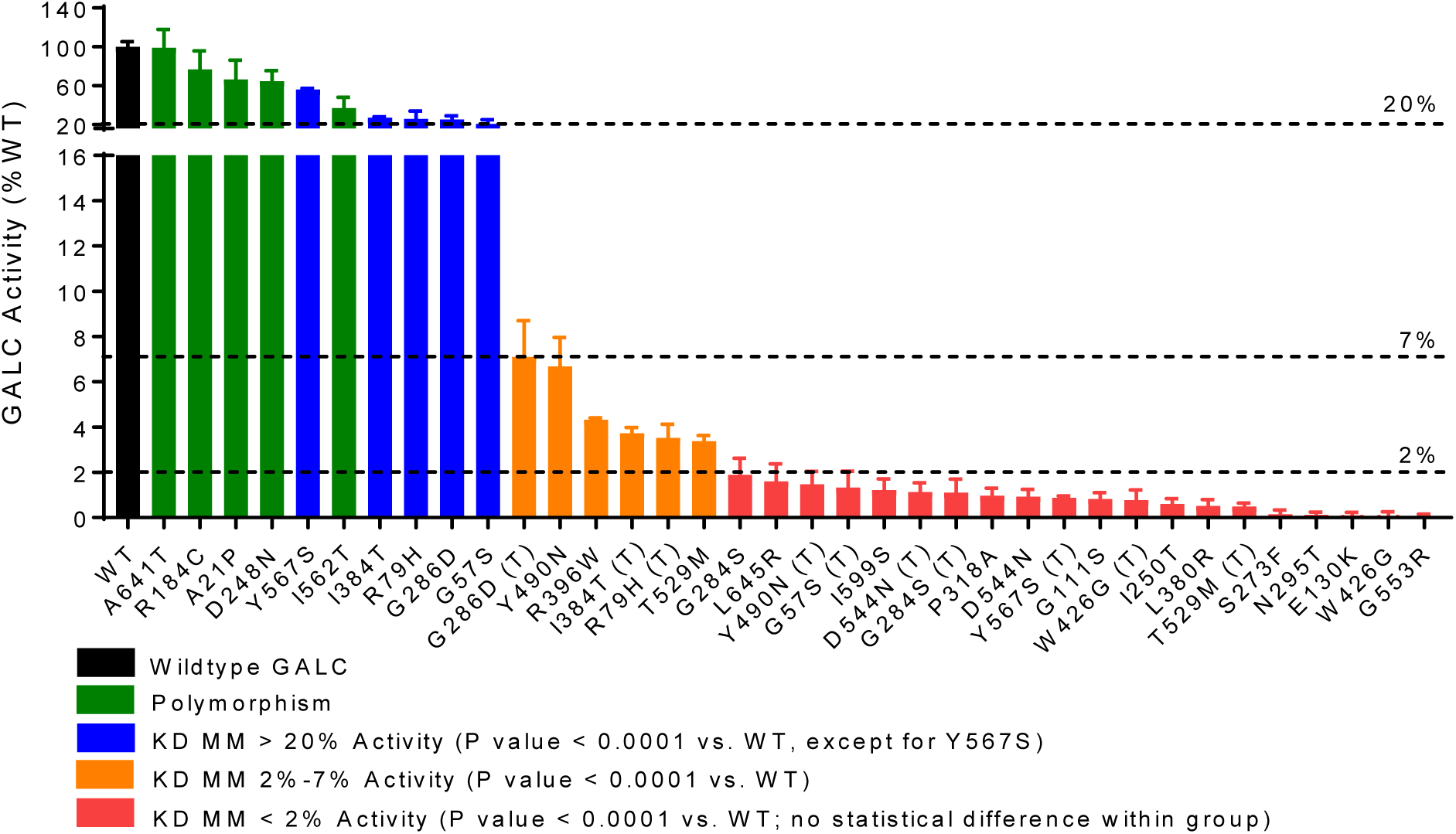
Dramatic reduction of GALC activity in 26 clinically-relevant MMVs. GALC activity in transiently-expressed cell models was measured in soluble cell lysate and presented as relative percent of WT-GALC. MMVs are ranked in descending order based on GALC activity. Of the MMVs showing significant reductions compared to WT-GALC, 6 variants had activity that remained relatively high (20-60% of WT, blue bars and I562T); 6 variants had low level activity (2-7% of WT, orange bars); and 20 variants had little to no activity (0-2% of WT, red bars). MMVs on the p.I562T background are annotated with (T). Data are presented as mean ± standard error of the mean from four independent experiments.

### GALC activity in MMV cell models positively correlates with age of symptom onset in KD patients carrying compound heterozygous or homozygous MMV genotypes

GALC MMVs are inherited as compound heterozygous or homozygous genotypes in KD patients. Here, we compared the age of symptom onset for published KD cases that contain a compound heterozygous MM-null mutation against the GALC activity observed in our MO3.13/ *GALC*-KO cells. Patients with these genotypes have MMVs for the p.I562T co-variant or an MMV on one allele and a null mutation, such as the 30 kb deletion, on the second allele (Figure 6A, “Het null”). MMVs that are associated with compound heterozygous genotypes include p.G553R, p.Y490N(T), p.I250T, p.S273F, p.P318A, p.G57S(T), p.T529M, p.Y567S(T), p.L645R, p.G248S, p.I384T(T) and p.G286D(T), shown in ascending age-of-onset order. Remarkably, GALC activity of these variants was highly correlated with the clinical severity observed in KD patients (Figure 6B, Pearson r = 0.94, P < 0.0001). For example, the most severe mutation, p.G553R, had undetectable GALC activity in our model and an average age-of-onset at 4.5 months. The least severe mutation, p.G286D(T), which maintained 7.1% of WT GALC activity, has an average age-of-onset at 60 months (juvenile-onset). In fact, all MMVs known to cause infantile KD (age-of-onset ≤ 12 months) had an activity less than 2% of WT in our model. By contrast, another variant known to cause juvenile-onset disease (p.I384T(T), age-of-onset = 42 months) had an activity of 3.7% (Figure 6A).

**Figure 6.**
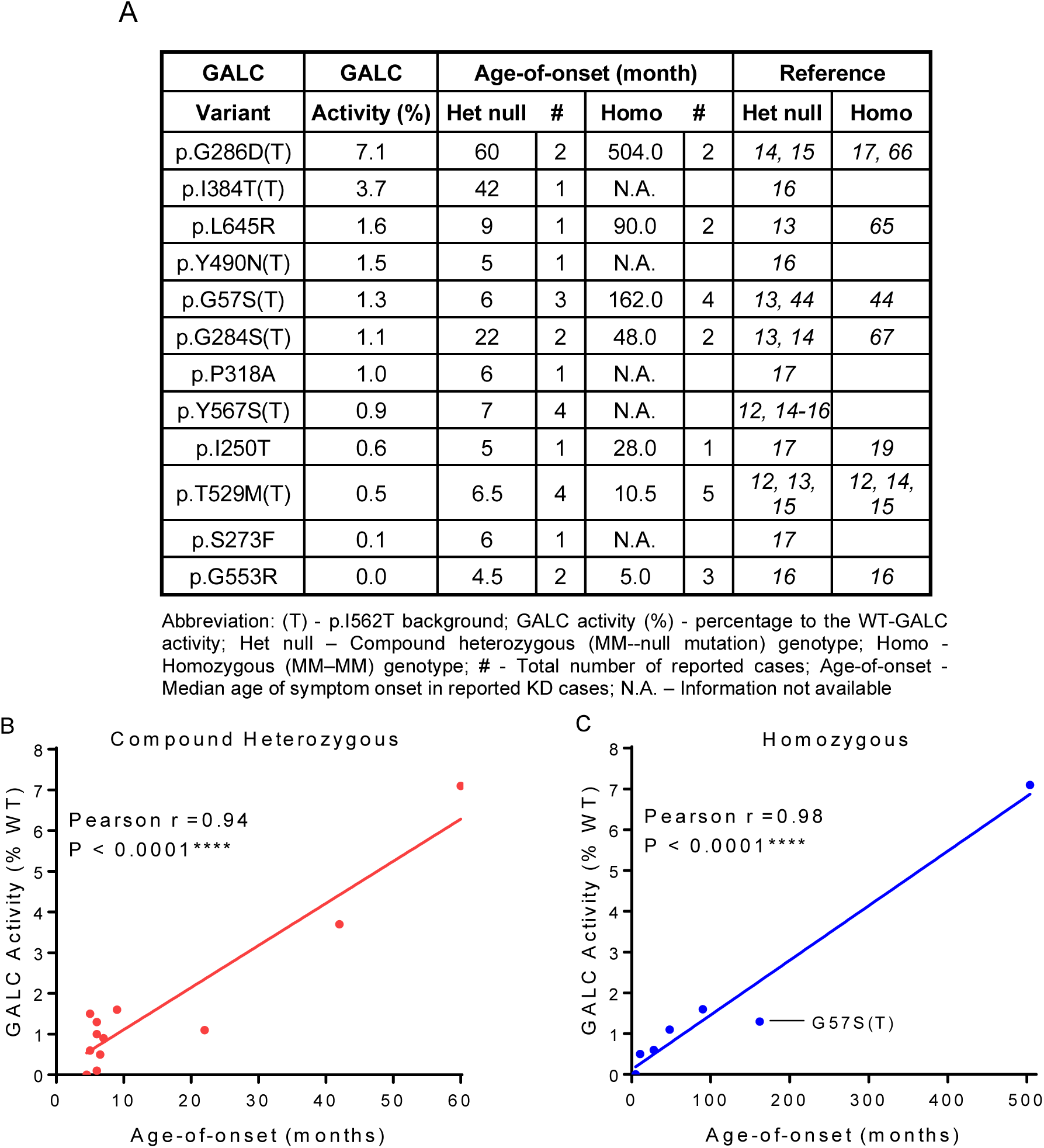
GALC activity in MMV cell models is highly correlated with the age-of-onset of clinical symptoms in KD patients carrying the same GALC MMV genotypes. (A) GALC activity levels in MMV cell models and the symptoms age-of-onset in KD patients with either compound heterozygous MM-null mutation or homozygous MM-MM genotypes, based on reported cases from literature. (B and C) Correlation analysis between GALC activity and the symptoms age-of-onset in KD patients with (B) heterozygous MM-null mutation genotypes (Pearson r = 0.94, P < 0.0001, n = 12) or (C) homozygous MM-MM genotypes (Pearson r = 0.98, P < 0.0001, n = 7).

After excluding cases that underwent treatment intervention, only 7 KD cases with homozygous MM-MM genotypes have been reported in published literature with an age-of-onset. Homozygous GALC MMVs associated with KD include p.G553R, p.T529M(T), p.I250T, p.G284S(T), p.L645R, p.G57S(T) and p.G286D(T), shown in ascending age-of-onset order (Figure 6A, “Homo”). Age-of-onset is consistently delayed (i.e. milder clinical severity) in patients with homozygous genotypes compared to patients with compound heterozygous MMVs. In part, due to higher residual GALC activity. For example, patients with homozygous p.I250T and p.L645R experience an average age-of-onset at 28 months and 90 months, respectively. By contrast, patients with the same compound heterozygous MMVs present with symptoms at 5 months and 9 months, respectively. p.G553R and p.G286D(T), inherited in the homozygous state, had the lowest and the highest activity in our cell models (0% vs. 7.1% of WT, respectively); which also translated into the earliest (5 months) and the latest (504 months) symptoms age-of-onset (Figure 6A). Strikingly, the activity of these variants was almost perfectly correlated with the age of symptom onset in KD patients (Figure 6C, Pearson r = 0.98, P<0.00001). An apparent outlier was the homozygous p.G57S(T) genotype, which was previous reported as a variant associated with longer survival^44^. Overall, we detect a strong genotype-phenotype correlation when comparing GALC activity in our MMV cell models and symptoms age-of-onset in KD patients with the *GALC* genotypes examined in our study. These results suggest that clinical onset in patients carrying these MMV genotypes may be predicted from residual GALC activity observed in the cell models and assays reported in this study.

### Lys-GALC protein levels correlate with GALC activity

Herein, we also analyzed the levels of lys-GALC protein in cells expressing the GALC MMVs panel. The cleaved, mature GALC fragment of WT-GALC can be detected as a 26 kDa lys-GALC protein band by the CL13.1 antibody, which targets the C-terminal fragment of GALC (Figure 7A-D). The p.D248N and p.I562T polymorphic variants reduced lys-GALC levels by 64% and 73%, respectively, compared to WT; whereas the pathogenic p.E130K and p.I250T variants reduced lys-GALC levels by 95% and 89%, respectively (Figure 7A, Table 3). Cells expressing the p.I562T co-variant tended to lower lys-GALC levels compared with the corresponding single MMV (Figure 7C, 7D and Table 3). Importantly, lys-GALC levels were positively and highly correlated to GALC activity among WT and all 36 MMVs examined (Figure 7E-7F, Pearson r = 0.93, P<0.0001). The strong correlation between activity and lys-GALC levels indicates that cleaved, mature GALC is the primary contributor to its activity. GALC-MMVs with less than 2% of WT activity, including p.G57S(T), p.R79H(T), p.G111S, p.S273F, p.N295T, p.L380R, p.I384T(T), p.T529M(T), p.G553R, p.Y567S(T), p.I599S and p.L645R, were largely undetectable by western blot (Table 3). These results further support the notion that deficiencies to GALC activity caused by GALC-MMVs is primarily due to reductions in lys-GALC levels.

**Figure 7.**
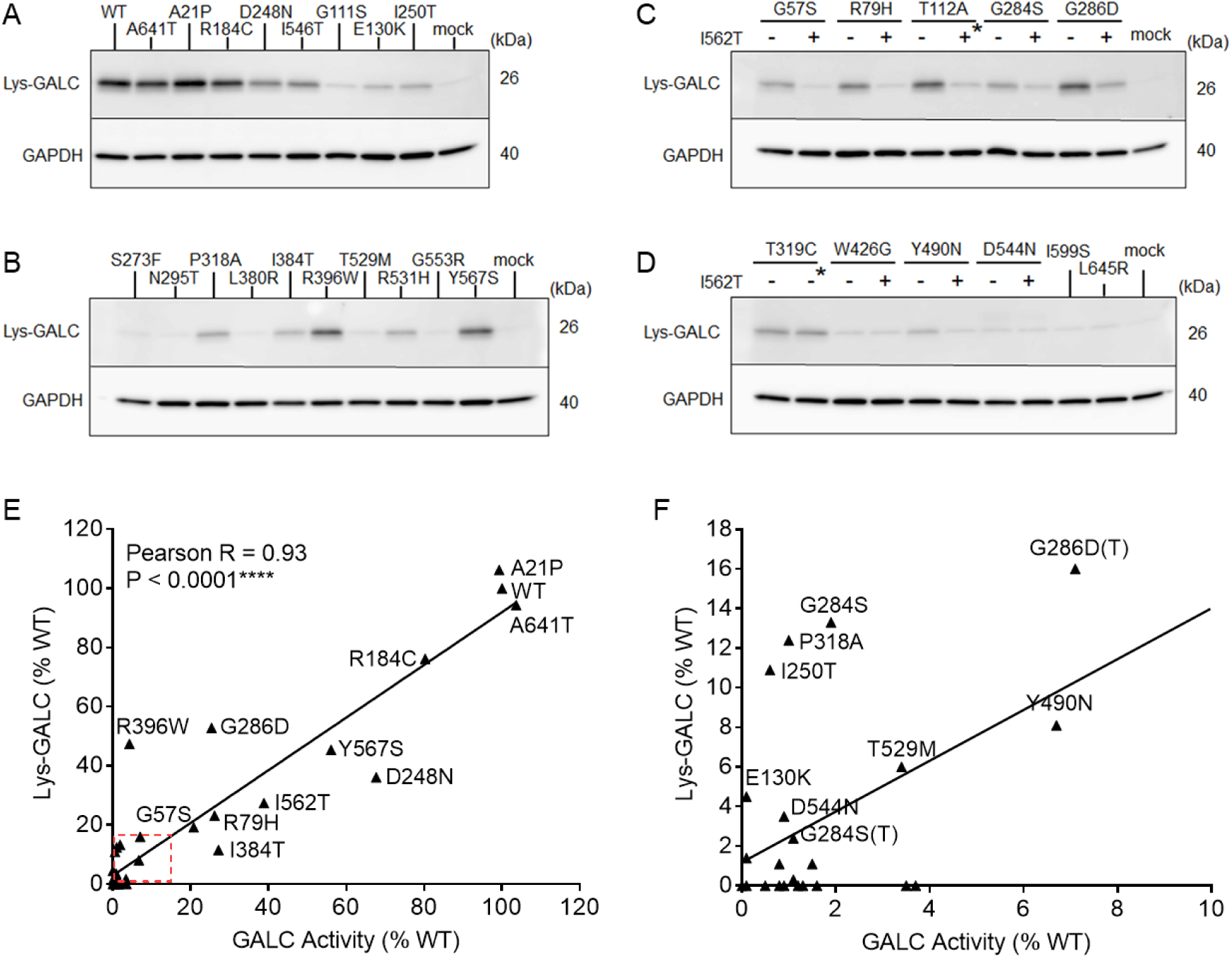
Lys-GALC levels are highly correlated with GALC activity. (A-D) Western blot results of lys-GALC and GAPDH in the lysate of GALC MMV expressing cell models. (C-D) show MMV in the absence (-) and presence (+) of the p.I562T polymorphic background except for 2 variants not under the scope of this study (asterisk). (E) Lys-GALC levels (normalized to GAPDH levels) are highly correlated with GALC activity in the GALC MMV cell models (Pearson r = 0.93, P < 0.0001, n = 37). (F) Expanded plot of the red line area shown in (E). MMVs with lys-GALC levels greater than 2% are included in (E) and (F).

### KD-related MMVs variably reduce secretion of GALC

To examine the effect of KD-related MMVs on protein trafficking, sec-GALC levels of MMVs were quantified by an in-house sandwich ELISA. Given that the carboxyl-terminal end of GALC is trimmed during the maturation process in lysosome, our sandwich ELISA captures the V5 tag on the C-terminal end of sec-GALC, followed by detection with a highly specific antibody against an internal epitope on GALC (CL13.1). In our cell mode, sec-GALC was measured at 8.6 ng/ml after WT-GALC was expressed for 72 hours. Polymorphic variants p.R184C and p.I562T reduced sec-GALC levels by 21% and 58%, respectively (Table 3). Eighteen of the 26 single MMVs (including polymorphic variants) also reduced sec-GALC levels (21% to 100%). Notably, 6 MMVs located on the β-sandwich domain or lectin domain of GALC (p.L380R, p.W426G, p.T529M, p.G553R, p.I599S and p.L645R) reduced sec-GALC more than 95% (Figure 8A). Additional p.I562T co-variants, including p.G57S(T), p.R79H(T), p.I384T(T), p.W426G(T), p.Y490N(T), p.T529M(T) and p.Y567S(T), also reduced sec-GALC by more than 95% (Table 3). Combined with our data that shows these 13 MMVs ameliorated GALC activity and lowered lys-GALC to undetectable levels, our results suggest that the observed GALC deficiency in these MMVs is likely via a mis-trafficking mechanism. The reduction of sec-GALC caused by individual MMVs coincides to the structural domain that contains the mutation. For example, 5 of 10 single MMVs located on the TIM barrel domain (57-353 aa) had no significant effect on sec-GALC levels. Alternatively, 9 of 11 single MMVs located on a carboxyl domain (354-684aa) reduced sec-GALC by at least 70% (Figure 8A). p.G111S, p.E130K, p.I250T, p.S273F and p.G286D, which are all found on the TIM barrel domain and in relatively close proximity to the active site^24^ (Figure 2), selectively reduced GALC activity and lys-GALC levels (Table 3). Taken together, our data suggests that these MMVs might selectively disrupt GALC trafficking to lysosome, but not to the extracellular space. Significant correlations between sec-GALC levels and age of symptom onset were not detected (Figure 8B and 8C). Therefore, we conclude that reduced GALC secretion is not a major factor contributing to clinical onset in KD.

**Figure 8.**
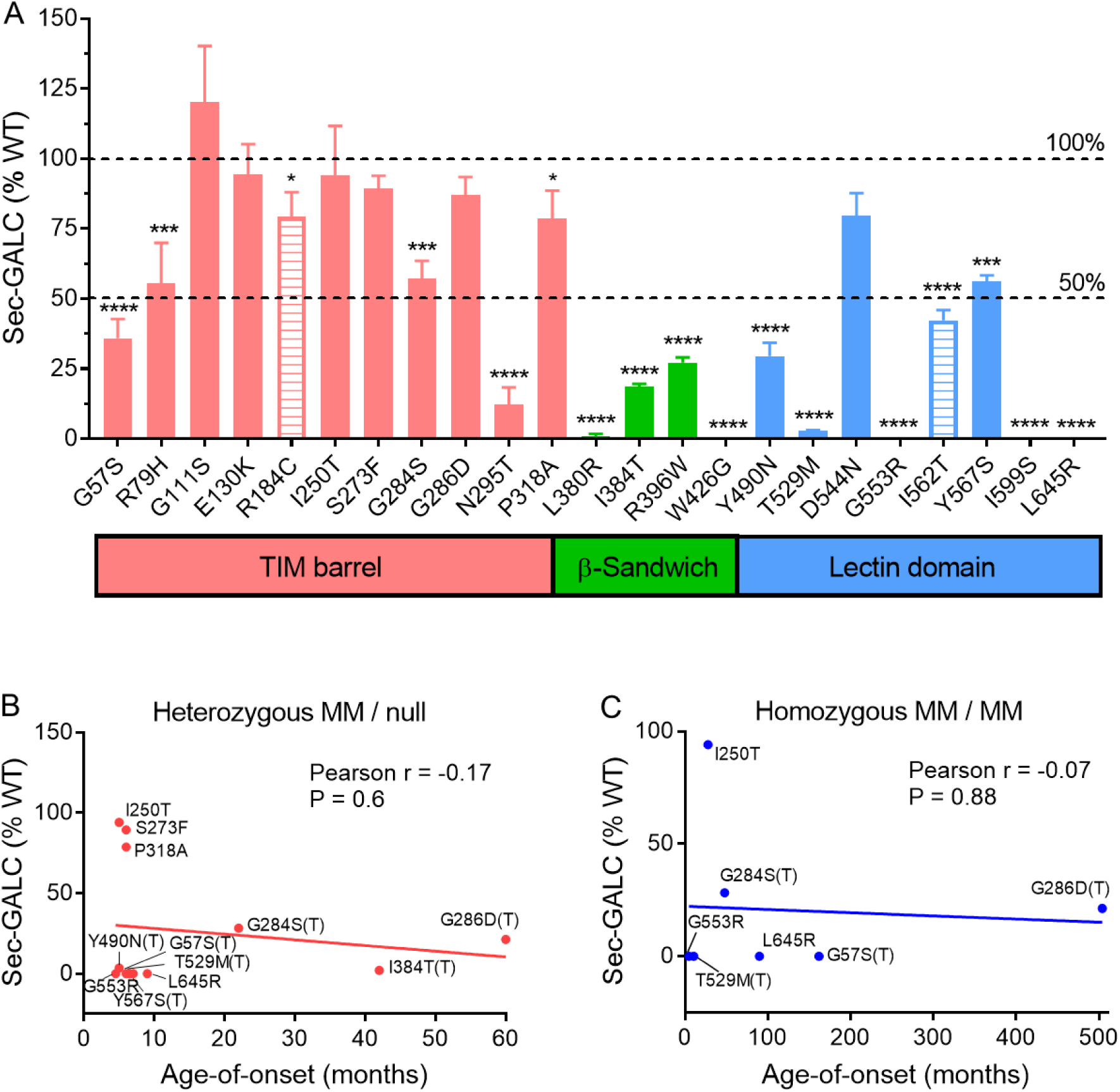
GALC secretion is more impaired by MMVs located on the β-sandwich and lectin protein domains but does not correlate to the onset of clinical symptoms in KD patients. (A) Sec-GALC levels are plotted against the structural location of GALC MMVs, categorized by their position on the TIM barrel (red), β-sandwich (green) and lectin (blue) domains. Polymorphisms are indicated by a striped bar. MMVs (11/12) significantly reduced sec-GALC levels compared with WT-GALC, specifically on the β-sandwich and lectin domains in our cell models. (Unpaired t-test, n=4, 95% confidence interval; * P < 0.05, ** P < 0.01, *** P < 0.001, **** P < 0.0001). (B, C) Analysis reveals no significant correlation between sec-GALC levels and the symptoms age-of-onset in KD patients with (B) heterozygous MM-null mutation genotypes or (C) homozygous MM-MM genotypes.

### KD MMs variably induce accumulation of intracellular pre-GALC

Intracellular retention of MM carrying proteins is a common pathogenic phenotype in many lysosomal storage disorders (LSDs), due in part, to the failure of endoplasmic reticulum-associated degradation (ERAD), which normally eliminates misfolded proteins over time^45, 46^. We measured retained, intracellular pre-GALC levels in transfected cells by ELISA using detergent-soluble cell lysate. The pre-GALC ELISA was validated by comparison to WB, which, as expected, revealed that the pre-GALC ELISA signal was positively correlated to the density of the 75 kDa pre-GALC protein band (Figure 9A, 9B, Pearson r = 0.71, P<0.0001, n = 35). Pre-GALC levels were elevated in cells expressing 10 MMVs and reduced with the expression of 2 MMVs (Figure 9C). In one case, the p.N295T variant, we observed a 6-fold increase in pre-GALC levels. This variant has previously been shown to induce abnormal glycosylation in cells^27^. Other MMVs (5) that caused elevated pre-GALC levels were also associated with a reduction in sec-GALC, including p.R184C (+54% pre-GALC; −21% sec-GALC), p.G57S (+99%, −64%), p.N295T (+510%, −88%), p.I384T (+89%, −70%) and p.Y490N (+67%, −78%). This data suggests that intracellular retention of GALC reduced secretion in these MMVs expressing cells (Table 3). Conversely, we identified a group of MMVs (p.L380R, p.W426G, p.T529M, p.G553R, p.I599S and p.L645R) that also reduced levels of sec-GALC, but did not increase pre-GALC levels; thus, the secretion defect in these variants is independent of intracellular retention.

**Figure 9.**
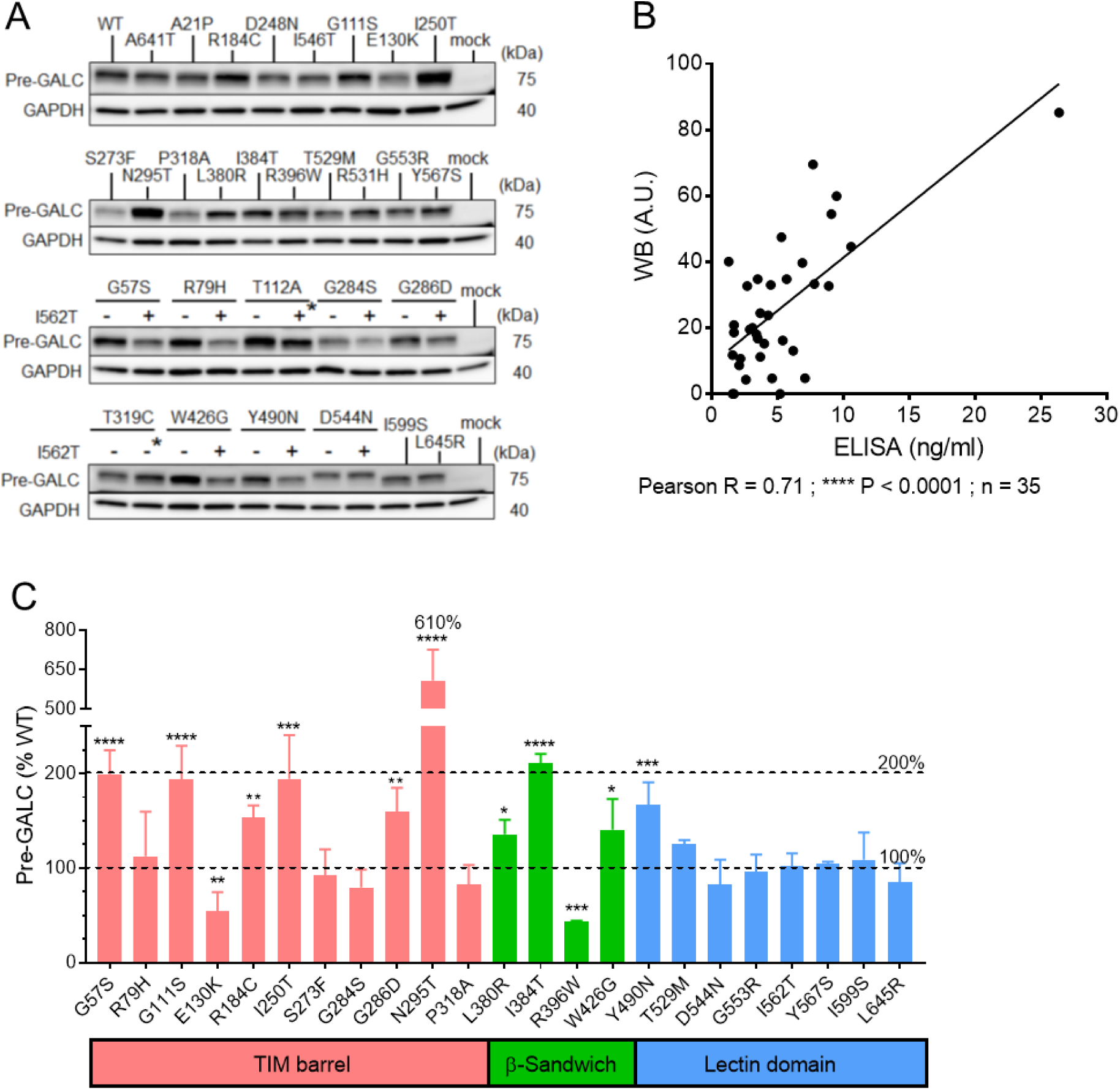
Accumulation of pre-GALC protein in GALC MMV cell models. (A) Intracellular pre-GALC and GAPDH proteins were detected by Western blot. The bottom 2 blots showed MMVs in the absence (-) and presence (+) of the p.I562T polymorphic background, with an asterisk indicating MMVs not under the scope of this study. (B) Pre-GALC levels (normalized to GAPDH) detected by Western blot are significantly correlated with pre-GALC levels measured by sandwich ELISA in our MMV cell models (Pearson r = 0.71, P < 0.0001, n = 35). (C) Pre-GALC levels measured by ELISA plotted against the structural location of the MMVs, categorized by their position on the TIM barrel (red), β-sandwich (green) and lectin (blue) domains. Ten out of 22 MMVs significantly increase pre-GALC levels compared with WT-GALC in the MMV cell models. (Unpaired t-test, n=3, 95% confidence interval; ** P < 0.01, *** P < 0.001, **** P < 0.0001).

### The p.I562T background synergistically impairs GALC function to reach pathogenic level

p.I562T is a known pseudo-deficiency variant associated with reduced levels of GALC activity. In fact, it was present in 87% of the 348 referral samples in the New York State Newborn Screen (NBS) Program after screening about 2 million samples^31^. Not only is it associated with pseudo-deficiency, this variant also frequently identifies as an allelic background with other MMVs on the *GALC* gene in confirmed KD cases^16, 47^. To evaluate the molecular effect of the p.I562T background on GALC protein, we analyzed the expression of a series of GALC p.I562T co-variants, including the p.G57S(T), p.R79H(T), p.G284S(T), p.G286D(T), p.I384T(T), p.W426G(T), p.Y490N(T), p.T529M(T), p.D544N(T) and p.Y567S(T) (Fig. 10). Expression of p.I562T alone greatly reduced WT-GALC activity (100% to 39%) in our cell model. Correspondingly, p.I562T co-expression synergistically reduced GALC activity in most variants (7 of 10); specifically, these variants, p.G57S (21% to 1.3%), p.R79H (26% to 3.5%), p.G286D (25% to 7.1%), p.I384T (27% to 3.7%), p.Y490N (6.7% to 1.5%), p.T529M (.4% to 0.5%) and p.Y567S (56% to 0.9%) (Fig. 10A). In contrast, p.I562T co-expression did not lower the activity observed with the p.G284S, p.W426G and p.D544N variants, which were already at very low activity (≤ 2%) when expressed alone (Table 3). These results suggest that the presence of the p.I562T allelic background can impair GALC function enough to reach the threshold of disease onset for these 7 MMVs.

**Figure 10.**
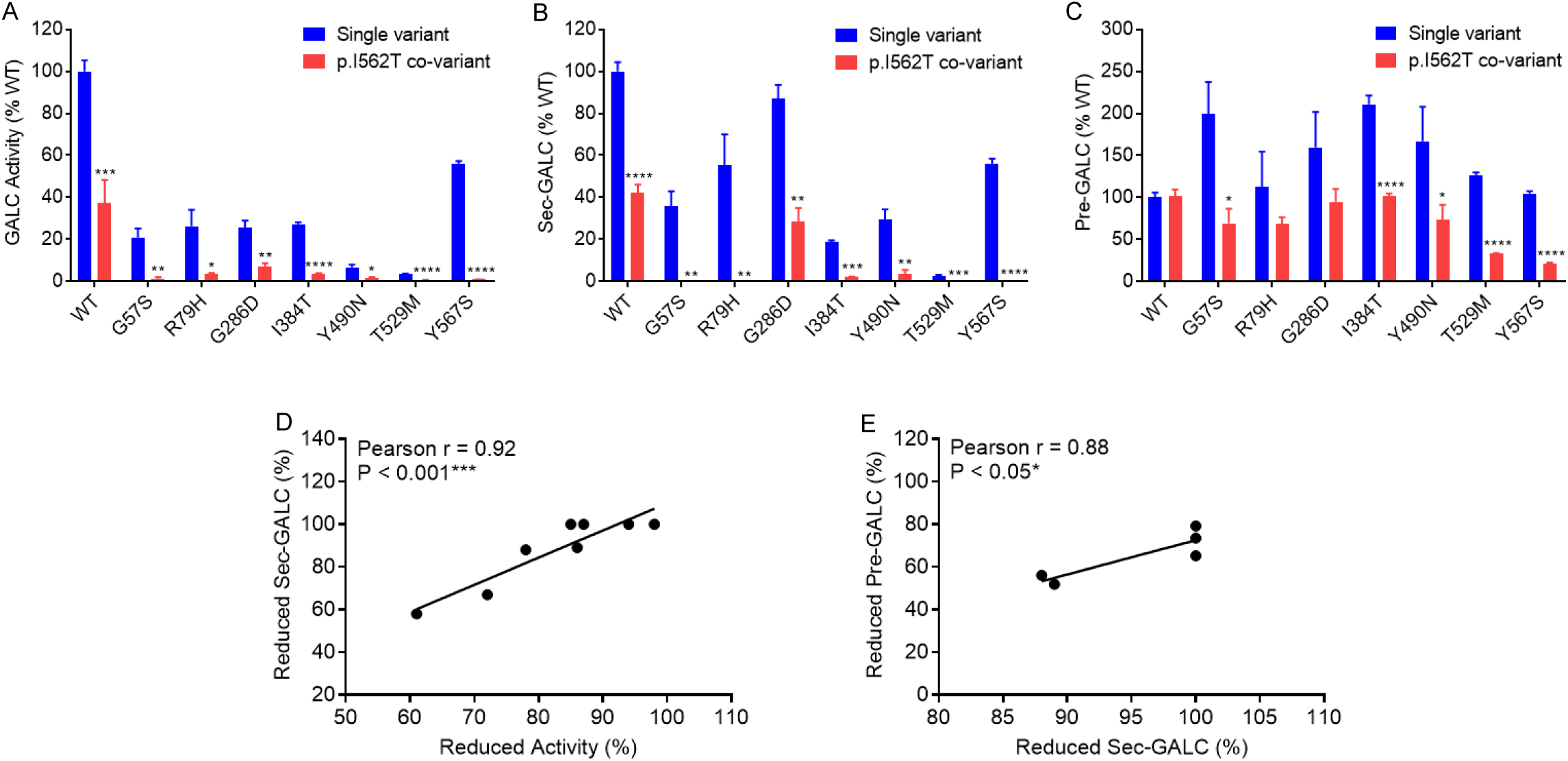
p.I562T background reduced GALC activity, sec-GALC levels and pre-GALC levels in the MMV cell models. (A) GALC activity, (B) sec-GALC levels and (C) pre-GALC levels from MMV expressing cells containing the p.I562T variant and seven other p.I562T co-variants (red bars) are compared side-by-side with those from WT cells or cells expressing the corresponding single variants (blue bars), respectively. (Unpaired t-test, n=4, 95% confidence interval; ** P < 0.01, *** P < 0.001, **** P < 0.0001). (B) The percentage reductions of GALC activity are highly correlated with the percentage reductions in sec-GALC levels in MMV expressing cells containing the p.I562T and seven other p.I562T co-variants (Pearson r = 0.92, P < 0.001, n = 8). (C) Reductions in sec-GALC levels (%)correlated with the reductions in intracellular pre-GALC levels (%) in five p.I562T co-variants (Pearson r = 0.88, P < 0.05, n = 5).

Interestingly, the p.I562T co-variants that resulted in reduced GALC activity, also caused a significant reduction in GALC secretion (Fig. 10B), and the two were highly correlated (Fig. 10D, Pearson r = 0.92, P < 0.001, n=8).This data suggests that the p.I562T background proportionally reduces the levels of correctly-folded GALC that gets trafficked to the both extracellular space and lysosomes. Mechanistically, p.I562T likely destabilizes the protein structure and induces the formation of misfolded GALC, which ultimately leads to an increase in degradation via protein quality control machinery. Indeed, intracellular pre-GALC levels were lower among most p.I562T co-variants compared to the corresponding single variants, although not all reached statistical significance (Fig. 10C). Among co-variants that statistically lowered pre-GALC levels (5 of 7), pre-GALC levels correlated to the reduced sec-GALC levels (Person r = 0.88, P < 0.05, n = 5); suggesting that p.I562T impairs GALC function through increased degradation of pre-GALC (Fig. 10E).

### GALC activity correlates with GALC secretion in the MMV cell models

Overall, we performed an expression study on WT-GALC and 36 GALC MMVs in the MO3.13/ *GALC*-KO cells to examine the effect of clinically-relevant variants on the residual activity, secretion and intracellular levels of GALC protein. Most MMVs we analyzed reduced GALC activity by more than 93% (26 of 36). Of these MMVs, half also reduced sec-GALC by more than 95%, and thus are likely to be associated with GALC insecretion – a protein mis-trafficking phenotype. Indeed, there was a positive correlation between GALC activity and sec-GALC level among all MMVs (Figure 11A, Pearson r = 0.5, P<0.01, n = 37). The moderate correlation was largely driven by single MMVs located on the TIM barrel domain, which elicited a unique low GALC activity/ normal sec-GALC profile. On the other hand, we did not detect any correlations between intracellular pre-GALC level and GALC activity (Figure 11B, Pearson r = −0.02, n = 37), or between pre-GALC and sec-GALC levels (Figure 11C, Pearson r = 0.01, n = 37). Taken together, this data suggests that among these variants, intracellular retention or degradation of pre-GALC is not a major determinant of the amount of GALC being trafficked to lysosomes or into the extracellular space.

**Figure 11.**
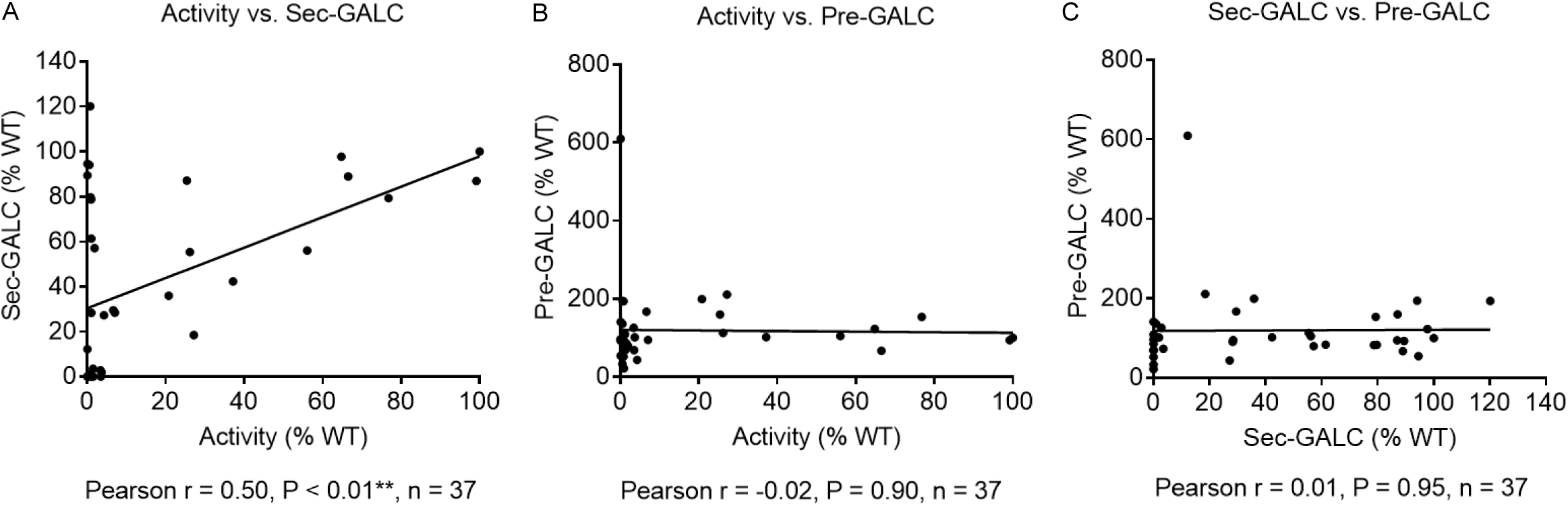
GALC activity correlates with sec-GALC levels in the MMV cell models. Correlations among (A) GALC activity, (B) sec-GALC levels and (C) pre-GALC protein levels in the GALC MMV expressing cells are analyzed using the Pearson correlation method. (A) GALC activity is significantly correlated with sec-GALC levels in the cell models (Pearson r = 0.5, P < 0.01, n = 37). No significant correlations are found between (B) GALC activity and pre-GALC levels, and between (C) sec-GALC and pre-GALC levels.

### GALC activity of MMVs correlates with psychosine levels in MMV cell models

Psychosine is a bona fide natural substrate of GALC, and psychosine levels reliably reflect the in-situ activity of GALC. Moreover, quantitative psychosine analysis was recently used to differentiate samples from KD patients with different clinical onset^48^. It is now well-accepted in the NBS setting that 10 nmol/L of psychosine detected in a dried blood spot is the cutoff value for infantile-onset KD cases that require referral for treatment intervention^49, 50^. Cell models of psychosine accumulation have been lacking, due to a general low abundance of endogenous psychosine in most cell types. To our knowledge, the MO3.13/ *GALC*-KO cell line used in this study is one of the few human cell models with a psychosine accumulation phenotype. We report a 21-fold increase in psychosine production in the *GALC*-KO cell line compared to parent cells (Figure 3E), which resembles the change in psychosine observed in brain tissue^5^. This robust readout cell model gives us the opportunity to examine the effect of pathogenic MMVs on psychosine accumulation in human cells.

To understand the correlation between clinical forms of KD and psychosine levels in our cell model, we performed a separate expression study on a subset of MMVs with published data for symptoms age-of-onset. Empty vector (mock), WT-GALC, p.I562T and 12 pathogenic MMVs were transfected into cells for 3 days. Transfected cells and medium were harvested and analyzed for cellular psychosine, residual GALC activity and sec-GALC (Table 4). At baseline, psychosine levels were high due to the absence of GALC. Transient transfection of WT-GALC led to expression of intracellular GALC in a sub-population of cells. However, GALC secreted from positively transfected cells was also taken up by neighboring non-transfected cells through a mechanism known as cross-correction. Therefore, psychosine was cleared by exogenous GALC regardless of transfection status of the cells. Transfection of WT-GALC reduced psychosine levels by 95% compared to mock control (from 0.348 to 0.016 pmol/mg). Notably, the difference in psychosine production (21-fold) found between *GALC-KO* and parent MO3.13 cells was identical to that observed between WT-GALC and mock transfected cells (Figure 3E), which suggests that the expression of WT-GALC normalized psychosine levels to match native cells. Transfection of the pseudo-deficiency variant p.I562T (46% GALC activity) and the late-onset variant p.G286D(T) (9% GALC activity), reduced psychosine levels by 82% and 73%, respectively, relative to mock control. The two most severe MMVs examined in this panel, p.T529M(T) (0.39% GALC activity) and p.G553R (0% GALC activity), only reduced psychosine levels by 20% and 2%, respectively. Affirmatively, there was a negative correlation between GALC activity and psychosine level among mock control, WT-GALC, p.I562T and the 12 pathogenic MMVs we analyzed (Figure 12A, Pearson r = −0.63, P<0.01, n = 15). These results confirm that GALC function impaired by pathogenic MMVs results in increased psychosine accumulation in our assay system.

**Table 4.** Psychosine level, GALC activity and sec-GALC level in MO3.13/ GALC-KO cells expressing a subset of clinically-relevant MMVs.

| GALC Variant | Psychosine |  |  | P value (vs.) |  | GALC Activity |  |  | P value (vs.) |  | Sec-GALC |  |  | P value (vs.) |  | Age-of-onset (month) |  |
| --- | --- | --- | --- | --- | --- | --- | --- | --- | --- | --- | --- | --- | --- | --- | --- | --- | --- |
|  | pmol/mg | SEM | % WT | WT | p.I562T | nmol /mg/hr | SEM | % WT | WT | p.I562T | ng/ml | SEM | % WT | WT | p.I562T | Het-null | Homo |
| Mock | 0.35 | 0.01 | 2154 |  |  | 0.00 | 0.00 | 0.00 |  |  | 0.00 | 0.00 | 0.00 |  |  |  |  |
| WT | 0.02 | 0.00 | 100 |  | **** | 5.45 | 0.60 | 100.00 |  | ** | 20.21 | 0.51 | 100.00 |  | *** |  |  |
| p.I562T | 0.06 | 0.00 | 399 | **** |  | 2.49 | 0.06 | 45.59 | ** |  | 11.69 | 0.58 | 57.85 | *** |  |  |  |
| p.G57S (T) | 0.19 | 0.01 | 1148 | **** | **** | 0.18 | 0.01 | 3.23 | *** | **** | 1.82 | 0.05 | 9.02 | **** | **** | 6 | 162 |
| p.I250T | 0.09 | 0.01 | 555 | ** |  | 0.07 | 0.01 | 1.22 | *** |  | 55.02 | 1.33 | 272.29 | **** |  | 5 | 28 |
| p.S273F | 0.25 | 0.02 | 1535 | *** |  | 0.03 | 0.01 | 0.49 | *** |  | 38.49 | 0.64 | 190.50 | **** |  | 6 | N.A. |
| p.G284S (T) | 0.13 | 0.01 | 800 | **** | *** | 0.09 | 0.01 | 1.66 | *** | **** | 9.52 | 0.17 | 47.11 | **** | * | 22 | 48 |
| p.G286D (T) | 0.09 | 0.01 | 585 | ** | * | 0.49 | 0.03 | 8.90 | ** | **** | 10.55 | 0.51 | 52.19 | *** | N.S. | 60 | 504 |
| p.P318A | 0.09 | 0.01 | 535 | *** |  | 0.11 | 0.02 | 2.09 | *** |  | 40.81 | 0.99 | 201.95 | **** |  | 6 | N.A. |
| p.I384T (T) | 0.13 | 0.01 | 827 | ** | ** | 0.28 | 0.01 | 5.15 | *** | **** | 0.76 | 0.10 | 3.74 | **** | **** | 42 | N.A. |
| p.Y490N (T) | 0.24 | 0.02 | 1491 | *** | *** | 0.03 | 0.01 | 0.60 | *** | **** | 1.56 | 0.12 | 7.74 | **** | **** | 5 | N.A. |
| p.T529M (T) | 0.28 | 0.01 | 1725 | **** | **** | 0.02 | 0.00 | 0.39 | *** | **** | 0.00 | 0.00 | 0.00 | **** | **** | 6.5 | 10.5 |
| p.G553R | 0.34 | 0.01 | 2112 | **** |  | 0.00 | 0.01 | 0.00 | *** |  | 0.00 | 0.00 | 0.00 | **** |  | 4.5 | 5 |
| p.Y567S (T) | 0.25 | 0.00 | 1184 | *** | *** | 0.09 | 0.02 | 1.70 | *** | **** | 0.00 | 0.00 | 0.00 | **** | **** | 7 | N.A. |
| p.L645R | 0.19 | 0.39 | 1339 | **** |  | 0.11 | 0.00 | 1.93 | *** |  | 0.00 | 0.00 | 0.00 | **** |  | 9 | 90 |

Statistical analysis is on subject data in comparison to WT-GALC expressing cells. MMVs on the p.I562T background were also statistically compared to p.I562T-GALC expressing cells. (Unpaired t-test, n = 3, 95% confidence interval, \*P < 0.05, \*\*P < 0.01, \*\*\*P < 0.001, \*\*\*\*P < 0.0001).
Abbreviations: pmol/mg – picomole per milligram total protein, SEM – standard error mean, Age-of-onset – age (months) that clinical symptoms appear in KD patients carrying the corresponding MMV in compound heterozygous or homozygous genotypes, Het-null – compound heterozygous MM/ null-mutation genotype, Homo – homozygous MM/MM genotype, N.S. – not significant, N.A. – information not available

**Figure 12.**
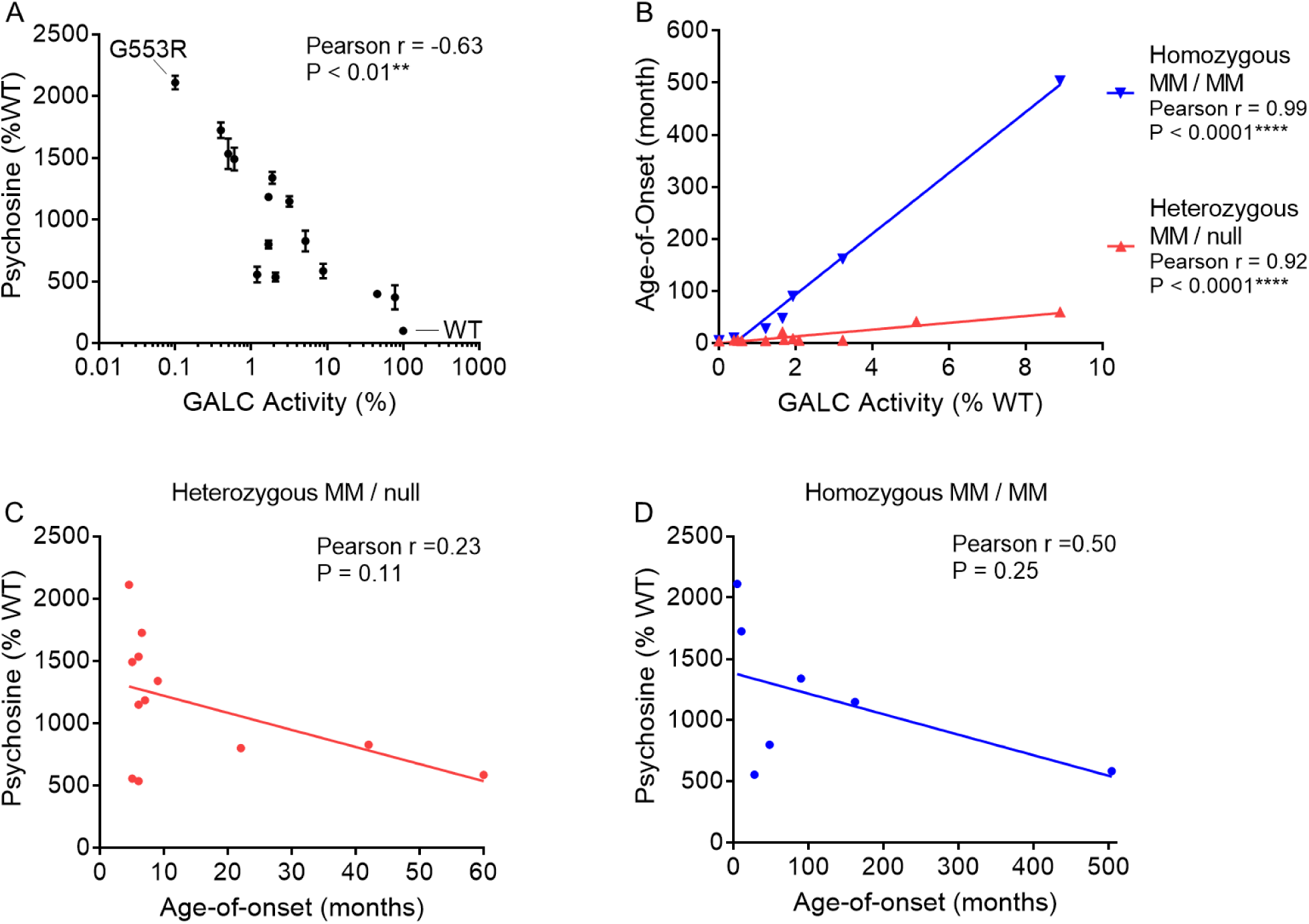
Cellular psychosine levels correlate with GALC activity in MMV cell models, but do not correlate with the onset of clinical symptoms in KD patients. (A) Expression studies were conducted in MO3.13/ *GALC*-KO cells for WT, p.I562T and 12 clinically-relevant GALC MMVs. Quantification of intracellular psychosine, GALC activity and sec-GALC levels are summarized in Table 4. Psychosine levels significantly correlated with GALC activity (Pearson r = −0.63, p < 0.01, n = 15). (B) Consistent with our transient expression study (Table 3, Figure 5), GALC activity was also highly correlated with the symptoms age-of-onset in KD patients with either homozygous MM-MM (Pearson r = 0.99, P <0.0001, n = 7) or compound heterozygous MM-null mutation genotypes (Pearson r = 0.92, P < 0.0001, n = 12). (C and D) However, psychosine levels were not significantly correlated with the age of clinical symptom onset, in either (C) compound heterozygous or (D) homozygous MMV genotypes.

Next, we performed correlation analysis between GALC activity and symptoms age-of-onset in KD cases with compound heterozygous and homozygous genotypes. Consistent with our results above (Figure 6), GALC activity observed in our model was highly correlated to the age-of-onset in both genotypes (Figure 12B, Pearson r = 0.92-0.99, P<0.0001). These results suggest that the results of our expression study, under our preset conditions, were consistent and reproducible. We then sought to understand whether psychosine levels in the cell model were correlated with clinical severity. Therefore, we performed correlation analysis between psychosine levels (relative to WT-GALC levels) and age-of-onset of KD cases with compound heterozygous or homozygous MMV genotypes. In many cases, psychosine levels in MMVs associated with infantile-onset KD, e.g. p.G553R and p.T529M(T) trended higher compared with that associated with late-onset KD (p.G286D(T)) (Table 4). Unfortunately, a significant correlation with either genotype was not observed (Figure 12C and 12D), indicating that psychosine levels in the cell models do not coincide with clinical age-of onset in KD.

## Discussion

In this study, we investigated the functional and trafficking effects of 21 pathogenic GALC MMVs and 10 additional MMVs found in cis with the pseudo-deficiency variant p.I562T in a *GALC*-KO human oligodendrocytic cell line. Our findings provide new insights into the molecular mechanisms that underly KD and highlight the potential for residual GALC activity measurements to inform patient diagnostics and prognostics.

### Impact of GALC MMVs on enzyme activity and clinical severity

Our results show that most GALC MMVs (20/31) dramatically reduce residual GALC activity, resulting in a ≥ 98% reduction in activity compared to WT-GALC. These results are consistent with the observation that clinically-relevant mutations in the *GALC* gene result in severe enzyme deficiencies in KD patients. Notably, MMVs found in infantile-onset patients with homozygous and compound heterozygous MM-null mutation genotypes, including p.S273F^17^, p.N295T^16^, p.P318A^17^, p.Y490N (T)^16^, p.G553R^16^, p.T529M (T)^13, 16^ and p.Y567S (T)^13, 16^, yielded < 2% of WT GALC activity in our model, which aligns with the early and severe clinical presentation seen in these patients. Conversely, cells expressing MMVs associated with later-onset forms of KD, such as p.G286D (T)^14, 16, 17^ and p.I384T (T)^16^, showed GALC activities between 2% and 7%, which justifies that observed in milder forms of the disease.

The strong correlation observed between residual GALC activity and age-of-onset in patients with compound heterozygous and homozygous MMV genotypes reinforces the notion that GALC activity obtained from this type of expression study in *GALC*-KO cells can serve as a reference for disease severity and as a potential tool for predicting the phenotype of a variant of uncertain significance (VUS) (Figure 6 and Figure 12B).

### Mechanisms of GALC dysfunction: Trafficking and maturation

Our study also sheds light on the mechanisms by which GALC MMVs affect enzyme function. We observed that many MMVs reduce levels of mature GALC (lys-GALC) in lysosomes, with a strong positive correlation between lys-GALC levels and GALC activity. This suggests that impaired trafficking of GALC to the lysosome is a key factor contributing to reduced enzyme activity. In addition, we observed many MMVs (13) that lead to dramatic reductions in both GALC secretion and lys-GALC levels for more than 96% (Figure 13). It indicates that these variants likely provoke misfolding and instability that affects protein trafficking to both extracellular space and lysosomes. In fact, MMVs, including p.L380R, p.W426G, p.G553R and p.L645R, that are expected to cause severe protein misfolding based on protein structure prediction^24^ are all associated with ultra-low GALC secretion (reduced > 99%), lys-GALC levels and activity (both reduced > 98%) in our study, highlighting the underlying mechanisms of GALC dysfunction.

**Figure 13.**
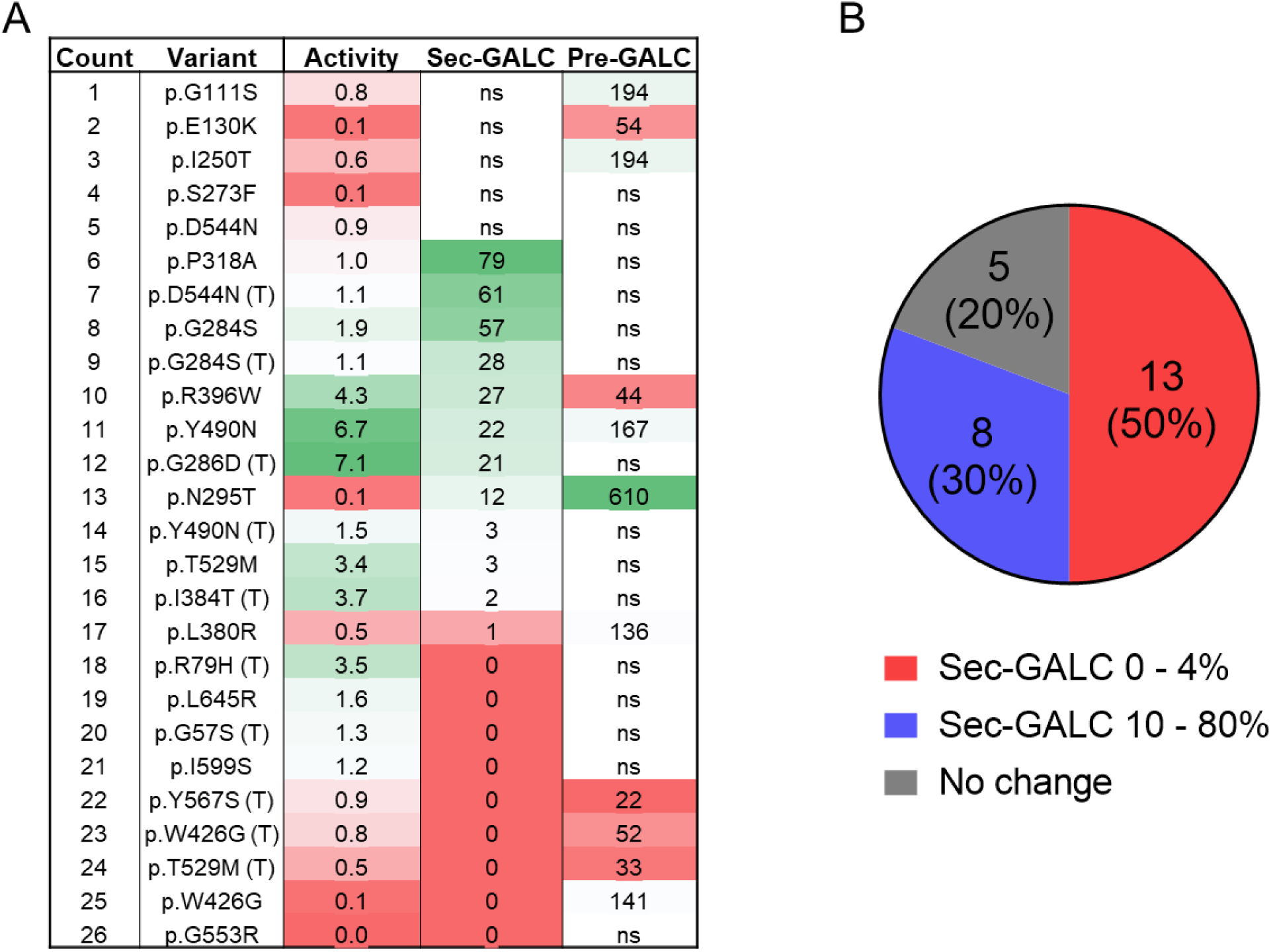
Dramatic reduction in GALC secretion in 50% of the low-activity GALC MMVs. (A) Summary table listing GALC activity, sec-GALC levels, and intracellular pre-GALC levels for 26 low-activity GALC MMVs. Values are presented as percentages relative to WT-GALC levels. The color spectrum from green to red indicates values from high to low within each group. “ns” denotes no significant difference from WT-GALC. (B) Distribution of sec-GALC levels among the 26 low-activity GALC MMVs.

Interestingly, 10 out of 36 MMVs also elevated levels of intracellular pre-GALC in our model, which suggests that these variants cause retention of the enzyme in the ER and/ or other intracellular compartments. The accumulation of pre-GALC could contribute to the observed deficiencies in enzyme secretion and activity. However, the fact that not all MMVs with reduced GALC activity caused an increase in pre-GALC levels, indicates that multiple mechanisms are likely at play, such as potential differences in protein stability and the trafficking pathway. p.G111S, p.E130K and p.I250T caused a dramatic reduction in GALC activity but not in secretion, which points to an alternative mechanism that selectively reduces lys-GALC function. Of relevance, an ARSA variant, p.P426L, a pathogenic MMV in metachromatic leukodystrophy, has been shown to selectively destabilize ARSA in lysosomes due to impairment of oligomerization^51^. It remains to be investigated whether the reductions to lys-GALC observed with these variants is related to a similar mechanism.

### Modifier effects of the p.I562T pseudo-deficiency variant

The presence of the p.I562T pseudo-deficiency variant significantly impacted the function of co-expressed GALC MMVs, leading to a synergistic impairment of GALC activity and secretion (Figure 10A and 10B). Given that many of these co-variants also elicited reductions in intracellular pre-GALC levels, we conclude that the p.I562T variant exacerbates the effects of other MMVs through a protein destabilization or expression inhibitory mechanism (Figure 10C). A previous study on physical properties of the p.I562T protein indicated that the polymorphism had only a subtle negative effect on its enzymatic activity and thermal stability compared to WT GALC. However, it had more impact on protein trafficking, because the variant could not be detected in the culture media of overexpressed HEK293T cells^27^. These findings are consistent with our results (50% reduction) (Table 3 and 4). Pseudo-deficiency variants were also identified in other LSDs, such as p.R247W and p.R249W in the *HEXA* gene ^52^, p.A300T in the *IDUA* gene ^53^, p.R595W in the *GLB1* gene ^54^, p.D152N in the *GUSB* gene ^55^ and p.D313Y in the *GLA* gene ^56^. Additionally, it has been reported that p.D313Y synergistically reduced α-galactosidase A function when expressed in cis with p.R112C or p.C172G, which were both identified in patients with classical Fabry disease^57^.

Given the high frequency with which p.I562T occurs in KD, cases of suspected KD sent for molecular testing should also undergo allelic DNA sequencing to reveal the p.I562T background status. Our results also highlight the need to include p.I562T background information in published literature and human genetic disease archives, such as the NCBI ClinVar and the Human Gene Mutation Database (HGMD), when reporting a variant as pathogenic or likely pathogenic. For example, both p.G286D and p.Y567S are classified as pathogenic or likely pathogenic on ClinVar. While GALC activity on both variants had below-threshold levels when expressed in cis with p.I562T (pG286D (T) at 7%, p.Y567S (T) at 1%), p.G286D and p.Y567S variants, when expressed alone, had GALC activity of 87% and 56% in our study, respectively (Table 3). Other studies consistently report high GALC activity for these 2 variants;^26, 30^ and both variants were uniformly found on the p.I562T background in all the low-GALC activity referral samples from the New York State NBS program^2^. Therefore, inheritance of either variant in its homozygous or compound heterozygous status is likely benign in the absence of p.I562T background^58^.

### Human MMV cell model using psychosine levels as a readout

Psychosine may be a more accurate readout than residual GALC activity for GALC function, as it has a broader dynamic range and is more stable in clinical samples^48^. Our results showed that it is feasible to establish human cell models with psychosine accumulation. Genetic knockout of *GALC* induced psychosine accumulation in the oligodendrocytic MO3.13 cell line. Functional expression of GALC MMVs by transient transfection catabolized psychosine accumulated in the KO cells in an activity-dependent manner. The measured residual GALC activity in the MMV cell models were significantly correlated to the psychosine levels (Figure 12A). However, the cellular psychosine levels did not correlate to the age of symptom onset in patients with these MMV genotypes (Figure 12C and 12D). The lack of clinical correlation with psychosine levels in the cell models is likely due to an insufficient penetrance of GALC expression in cell populations by transient transfection. In addition, efficiency of cross-correction by secreted GALC may be low or absent in MMVs that severely impact GALC secretion. Therefore, technical strategies that can increase penetrance, such as increasing transfection efficiency and/or positive selection of GALC expressing cells, will likely improve the psychosine-clinical phenotype correlation in cellular models.

### Implications for Clinical use and therapeutics

Genotype-phenotype correlations in KD have not been well established. In particular, the topic on functional deficiency caused by MMV genotypes has not been explored, although it is the most common type of pathogenic mutation. Our study highlights the importance of assessing residual GALC activity in combination with genetic testing to better understand the clinical implications of GALC MMVs. The correlation between residual GALC activity determined in the *GALC*-KO cell model and the symptoms age-of-onset provides a basis for using enzyme activity measurements as a predictive tool for clinical severity. Our plan is to expand the list of clinically-relevant MMVs as more clinical information becomes available, which would facilitate more accurate prognostic assessments and inform management strategies for KD patients with MMV genotypes.

Our study also identifies MMVs that likely impair GALC function via a protein misfolding mechanism. Pharmacological chaperones (PC) are small molecules that can act to restore protein folding by binding to the protein’s active site within the ER, thus stabilizing the misfolded proteins. 1-Deoxygalactonojirimycin, originally identified as a reversible, competitive inhibitor of α-galactosidase A (AGAL), was the first PC approved by the Food and Drug Administration to treat patients with Fabry disease^59^. For KD, PC candidates, such as iso-galacto-fagomine (IGF) have been shown to bind GALC at nanomolar concentrations and increase thermal stability of the protein ^60^. Our current cell model and assays would be an ideal *in vitro* platform to identify and develop therapeutics, such as PCs for KD. GALC activity and psychosine levels could be considered functional readouts to determine potency, efficacy and optimal dosing regimens. In addition, the platform would allow screening of mutations amenable to specific drugs, similar to the GLP-HEK assay used for Fabry disease^61^.

## Conclusion

In summary, our study provides a comprehensive analysis of the functional and trafficking effects of GALC MMVs in a *GALC*-KO cell model. The results offer valuable insights into the pathogenic mechanisms associated with the mutations and support the use of residual GALC activity as a tool to establish genotype-phenotype correlations in KD. Continued research into the molecular mechanisms of GALC dysfunction will be essential for advancing our understanding of KD biology and developing MMV-specific therapeutic interventions.

## Conflict of Interest Statement

Authors declare no conflict of interest.

## Funding

This work is partly funded by The Rosenau Family Research Foundation to C.W.L. (Investigator grant: 2022 to 2025) and the MidAtlantic Neonatology Associates research fund. The Vermont Biomedical Research Network Proteomics Facility (RRID: SCR_018667) is supported through NIH grant P20GM103449 from the IDeA (Institutional Development Award) Networks of Biomedical Research Excellence (INBRE) Program of the National Institute of General Medical Sciences.

## Supplemental Information

Materials and Methods: Proteomic analysis of native MO3.13 cells (WT) and *GALC*-KO cells (KO) using isobaric tandem mass tags

Supplemental Table 1. Primers for site-directed mutagenesis

## Notes

### Competing Interest Statement

The authors have declared no competing interest.

