## Supplemental Information for "Expression study of Krabbe Disease *GALC* missense variants – Insights from quantification profiles of residual enzyme activity, secretion and psychosine levels"

Materials and Methods: Proteomic analysis of native MO3.13 cells (WT) and *GALC*-KO cells (KO) using isobaric tandem mass tags

Supplementary Table 1: Primers for site-directed mutagenesis

### Proteomic analysis of native MO3.13 cells (WT) and *GALC*-KO cells (KO) using isobaric tandem mass tags

*Sample preparation and trypsin digestion.* Total protein (20 µg) from each sample was loaded onto the SDS-PAGE and run slightly (~ 1 cm) into the separating gel, which was stained with Coomassie Brilliant Blue. SDS-PAGE allowed assessment of protein loadings and removal of any incompatible detergents for subsequent in-gel trypsin digestion. Single wide bands containing briefly separated proteins from each individual sample were excised, minced to approximately 1 $mm^3$ cubes, followed by destaining with 50% acetonitrile ($CH_3CN$)/50 mM triethyl ammonium bicarbonate (TEAB). After dehydrating with $CH_3CN$, reduction and alkylation of disulfides were conducted with 10 mM dithiothreitol and 50 mM iodoacetamide in 100 mM TEAB, respectively. After repeated washing and dehydrating with 100 mM TEAB,and $CH_3CN$, the gel pieces were vacuum dried and swelled with trypsin solution (Promega V511A, 15 µg/ml in 40 mM TEAB, 5% $CH_3CN$) and incubated overnight at 37°C. Peptides were extracted successively with formic acid and acetonitrile then dried under vacuum.

*Peptide labeling by Tandem Mass Tags and high pH fractionation.* The labeling procedures were performed according to the manufacturers' protocols (Thermo Fisher Scientific, Waltham, MA, USA). Briefly, 10 µg of the dried peptides from each sample resuspended in 50µL of TEAB, and 0.8 mg of TMT reagents were warmed to room temperature and dissolved in 41 µL of anhydrous $CH_3CN$. 20.5 uL of the TMT reagents were added to respective peptide sample solutions, followed by briefly vortexing and an incubation for 1 h at room temperature with shaking. The labeling was verified by short mass spectrometry runs and database searches, and re-labeling is conducted to ensure that > 99% of the peptides were labeled in individual samples before combining. The reactions were then quenched by adding 8 µL of 5% hydroxylamine and combined. The labeled peptides were fractionated using the high-pH reversed-phase spin column (Cat. No.: 84868; Thermo Scientific) into 8 fractions by LC/MS. All samples were kept at -80 °C until mass spectrometry analysis.

*Liquid chromatography-tandem mass spectrometry (LC-MS/MS).* The fractionated TMT labeled peptides were resuspended in 2.5% $CH_3CN$ and 2.5% formic acid (FA) in water. One-fifth of the labeled peptides were analyzed. Mass spectrometry was performed on the Orbitrap Fusion mass spectrometer coupled to an EASY-nLC ULTRA (nLC-1000, Thermo Scientific, Waltham, MA, USA). Samples were loaded onto a 100 µm x 260 mm capillary column packed with UChrom C18 material (1.8-µm 120 Å Uchrom C18, nanoLCMS Solutions, CA, USA) at a flow rate of 300 nl $min^{-1}$. The column end was laser pulled to a ~3 µm orifice and packed with minimal amount of 5-um Magic C18AQ before packing with the 1.8-µm particle. Peptides were separated using a gradient of 5-7% $CH_3CN$/0.1% FA over 5 min, 7-32% $CH_3CN$/0.1% FA in 120 min, 32-90% $CH_3CN$/0.1% FA in 10 min then 90% $CH_3CN$/0.1% FA for 10 min, followed by an immediate return to 0% $CH_3CN$/0.1% FA and a hold at 0% $CH_3CN$/0.1% FA. Peptides were introduced into the mass spectrometer via a nanospray ionization source with a spray voltage of 2.2 kV. “Top speed in 2.5 seconds” acquisition mode and TMT specific Synchronous Precursor Selection (SPS) workflows were used to acquire mass spectrometry data. Survey scan from *m/z* 400-1600 at 120,000 resolution (AGC target $4e^5$ (100% normalized); max IT 50 ms; profile mode) was acquired with lock mass function activated (*m/z* 371.1012; use lock masses: best;

lock mass injection: full MS), followed by data-dependent collison-induced dissociation (CID) tandem mass spectrometry (MS/MS) scans on the most abundant ions (AGC target 1e$^{4}$ (standard); Max IT: Auto; Resolution: Turbo) in the ion trap with a normalized collision energy (NCE) at 35% and an isolation width of 0.7 *m/z*. MS3 was performed using SPS with isolation widths of 3 Da and an NCE of 65% (AGC target: Custom: 300%; Max IT: Auto). The product ions from MS3 were scanned from *m/z* 100 – 500 in the Orbitrap with a resolution of 50,000. Dynamic exclusion was enabled (peptide match: preferred; exclude isotopes: on; underfill ratio: 1%; exclusion duration: 60 sec; singly charged ion excluded). Advanced Peak Detection was set to "off". The samples were injected twice as technical replicates.

*Database searches.* The 16 mass-spectrometry .RAW files (8 high-pH reverse phase separation fractions analyzed in 2 technical replicates) generated from each experiment were imported into the Proteome Discoverer 2.4 (Thermo Fisher Scientific) as "fractions". Product ion spectra were searched using the SEQUEST in the Processing workflow against a Uniprot *Homo Sapiens* protein database (UP000005640; downloaded in July., 2019). Search parameters were as follows: (1) full trypsin enzymatic activity; (2) maximum missed cleavages = 2; (3) minimum peptide length = 6; (4) mass tolerance at 10 ppm for precursor ions and 0.6 Da for fragment ions; (5) dynamic modifications on methionines (+15.9949 Da: oxidation), dynamic modification on protein terminus (+42.01 Da: Acetyl; -131.040 Da: Met-loss; -89.030 Da: Met-loss+Acetyl) , static TMT6plex modification (The TMT6plex and TMT10plex have the same isobaric mass) on peptide N-termini and lysines (229.163 Da); (6) 4 maximum dynamic modifications allowed per peptide; and (7) static carbamidomethylation modification on cysteines (+57.021 Da). Percolator node was included in the workflow to limit the false discovery rates (FDR) to less than 1% in the data set.

*Quantification and statistical analysis.* The biological replicates and technical replicates were assigned accordingly in the “Study Factors” tab (WT_1, WT_2, WT_3, WT_4, KO_1, KO_2, KO_3, KO_4) and “Samples” tab, respectively in the Proteome Discoverer. The 8-plex workflows were constructed by omitting 130C and 131. The abundances of TMT labeled peptides were quantified with the Reporter Ions Quantifier node in the Consensus workflow and parameters were set as follows: (1) both unique and razor peptides were used for quantification; (2) Reject Quan Results with Missing Channels: False; (3) Apply Quan Value Corrections: True (values set according to the product spreadsheet (Lot #: VD296213); (4) Co-Isolation Threshold: 75; (5) Average Reporter S/N Threshold = 10; SPS Mass Matches [%] = 65; (6) Normalization mode: “total protein abundance”; and (7) Scaling Mode was set “on All Average”. Non-nested design was used for the analysis of biological replicates (two technical replicates for each of the 4 biological replicates). Protein ratio calculation was “Protein Abundance Based”. Two-tailed t-test was used for hypothesis testing and adjusted p-values were calculated by Benjamini-Hochberg procedure. All the protein identification and quantification information (<1% FDR; with protein grouping enabled) was exported from the Proteome Discoverer result files to Excel spreadsheets. Fold change ($\log_2$) and p values ($-\log_{10}$) information and scaled abundances (the abundances were scaled so that of proteins of varying abundances could be represented on the same color intensity scale of the heat map) were imported into Graph Pad Prism 8 (GraphPad Software Inc., CA) for constructing volcano plots and heat maps. The mass spectrometry / proteomics data, including raw (.raw), search (.msf), and analysis (.sf3) files, as well as MGF and mzIdentML files exported from Scaffold, have been deposited to the ProteomeXchange Consortium via the PRIDE partner repository with the dataset identifier.

**Supplemental Table 1**: Primer of site-directed mutagenesis

| GALC Variant | Forward primer | Reverse primer |
|---|---|---|
| p.A641T | GGGTCATTTCACCTCTGGCATGC | TTAATAGTTAACGTGAGTGTATACC |
| p.A21P | GACTGCGGCCCCGGGTTCGGC GG | ATAGCTTTCGCTCGGCGTTGCCAG |
| p.R184C | GGGCGCCAAGTGTTACCATGATT | ACAATCCAGGTCACGACATAATAG |
| p.D248N | CAAGGTGGTTAATGTTATAGG GG | AAGAGTTCGGCATCAAGG |
| p.I562T | CAA TCT GAC TAC AAA GTG TGA TGT ATA C | GTC CAG TTG TAG TCT CCT ATA ATA C |
| p.G57S | GGA GTT CGA CAG CAT CGG CGC GG | CGG CCC AGC CCG TCG GAG |
| p.R79H | AGA GCC CTA TCA TTC TCA GAT ATT G | GGG TAA TTT ACT AGA AGT CG |
| p.G111S | GAC AAC AGA CAG CAC TGA GCC CT | TGC CCA TCA CCA CCT ATT TC |
| p.E130K | CCG AGG ATA CAA GTG GTG GTT GA | AAA TAA TTC TCA TCT AGT GCA TAA TGC |
| p.I250T | GGT TGA TGT TAC AGG GGC TCA TT | ACC TTG AAG AGT TCG GCA |
| p.S273F | GCT TTG GTC TTT TGA AGA CTT TAG | TTC TTC CCA GTC AAC TTT G |
| p.G284S | TAG TGA CAT GAG TGC AGG CTG CT | TTT AAA GTG CTA AAG TCT TCA GAA GAC |
| p.G286D | CAT GGG TGC AGA CTG CTG GGG TC | TCA CTA TTT AAA GTG CTA AAG TCT TCA GAA GAC C |
| p.N295T | TTT AAA TCA GAC TTA TAT CAA TGG CTA TAT GAC TTC CAC AAT CGC | ATG CGA CCC CAG CAG CCT |
| p.P318A | TGA ACA GTT GGC TTA TGG GAG AT | TAG TAA CTA GCC ACT AAA TTC C |

| | | |
|---|---|---|
| p.L380R | CTT AGG GAA CCG CAC CAT CAT CA | CCA TCA GTC AGA GCT ACG |
| p.I384T | CAC CAT CAT CAC TGA AAC CAT GA | AGG TTC CCT AAG CCA TCA |
| p.R396W | TAA GTG CAT ATG GCC ATT TCT TC | GAA TGT TTA TGA CTC ATG GTT TC |
| p.W426G | GCT ACA GGT AGG GTA TAC CAA AC | TCT GGT ATT TCA CTA AAA GAT CC |
| p.Y490N | CCC AAG TAC CAA TAA GGA TGA TTT C | AAG GGC TGG GAT TTT GGA |
| p.T529M | GCA TCA CTT CAT GCT ACG CCA AG | TCG CCA GGG TCT TCA ATA TTT G |
| p.D544N | GTG GGC TGC CAA TGC ATC CAA CA | GTA ATG GGT CTC TGG TTG AGA AC |
| p.G553R | CAG TAT TAT AAG AGA CTA CAA CTG | ATT GTG TTG GAT GCA TCG |
| p.Y567S | GTG TGA TGT ATC CAT AGA GAC CC | TTT ATA GTC AGA TTG GTC C |
| p.I599S | TTT CTT CTG GAG TTT TGC AAA TG | ATT CCT CTG GCA CTT CTA ATC |
| p.L645R | CTC TGG CAT GCG GAA TGA CAA GT | GTG AAA TGA CCC TTA ATA GTT AAC |
